# Comparisons of dual isogenic human iPSC pairs identify functional alterations directly caused by an epilepsy associated *SCN1A* mutation

**DOI:** 10.1101/524835

**Authors:** Yunyao Xie, Nathan N. Ng, Olga S. Safrina, Carmen M. Ramos, Kevin C. Ess, Philip H. Schwartz, Martin A. Smith, Diane K. O’Dowd

## Abstract

Over 1250 mutations in *SCN1A*, the Nav1.1 voltage-gated sodium channel gene, are associated with seizure disorders including GEFS+. To evaluate how a specific mutation, independent of genetic background, causes seizure activity we generated two pairs of isogenic human iPSC lines by CRISPR/Cas9 gene editing. One pair is a control line from an unaffected sibling, and the mutated control carrying the GEFS+ K1270T *SCN1A* mutation. The second pair is a GEFS+ patient line with the K1270T mutation, and the corrected patient line. By comparing the electrophysiological properties in inhibitory and excitatory iPSC-derived neurons from these pairs, we found the K1270T mutation causes cell type-specific alterations in sodium current density and evoked firing, resulting in hyperactive neural networks. We also identified differences associated with genetic background and interaction between the mutation and genetic background. Comparisons within and between dual pairs of isogenic iPSC-derived neuronal cultures provide a novel platform for evaluating cellular mechanisms underlying a disease phenotype and for developing patient-specific anti-seizure therapies.

## 1. Introduction

Genetic epilepsy with febrile seizure plus (GEFS+) is a form of epilepsy with febrile seizures that persists beyond six years of age (Scheffer and Berkovic, 1997). The clinical phenotype of GEFS+ is characterized by febrile seizures beginning in infancy that persist beyond 6 years of age and in some cases individuals also experience myoclonic, atonic and/or absence seizures (Miller and Sotero de Menezes, 2014). Mutations in a variety of genes that encode ion channels and neurotransmitter receptors have been identified in individuals with GEFS+ (Baulac et al., 2001; Dibbens et al., 2004; Escayg et al., 2001, 2000; Harkin et al., 2002; Sugawara et al., 2001; Wallace et al., 2002, 2001a, 2001b, 1998). One hot spot for mutations causing GEFS+ is the *SCN1A* gene which encodes the alpha subunit of the Nav1.1 voltage-gated sodium channel. Approximately 50 missense mutations at locations throughout *SCN1A* are associated with different individuals diagnosed with GEFS+ (Meng et al., 2015). There are also multigenerational families in which individuals with the same *SCN1A* mutation exhibit a broad range of seizure phenotypes (Abou-Khalil et al., 2001; Goldberg-Stern et al., 2014).

The heterogeneity of disease phenotypes arising from different mutations in the same gene, and from the same mutation between individuals complicates understanding of the underlying cellular mechanisms. Model systems that aid in mapping the links between individual *SCN1A* mutations and their cellular effects can facilitate development of more effective patient-specific drug therapies. Zebrafish, knock-in *Drosophila* and mouse models have all been important tools in this endeavor (Schutte et al., 2016) but evolutionary distances leading to species-specific mechanisms may lead to failed therapies in patients (Marian, 2011; Maxwell et al., 2014).

In recent years, patient-derived induced pluripotent stem cells (iPSCs) that can be differentiated into neurons have emerged as a new model to explore how epilepsy-associated *SCN1A* mutations affect activity and sodium channel properties in human neurons. These studies have focused on mutations associated with Dravet syndrome (DS), a form of epilepsy that is typically more severe than GEFS+. Previous studies in iPSC-derived neurons have shown that nonsense R1645X, and missense S1328P and Q1923R mutations in *SCN1A* associated with DS confer impaired excitability and reduced sodium current density in inhibitory neurons, giving rise to hyperactivity in neural networks (Higurashi et al., 2013; Liu et al., 2016; Sun et al., 2016). However, examination of the DS-associated *SCN1A* missense mutations, Q1923R and F1415I, revealed increased excitability and increased sodium current amplitudes in excitatory neurons (Jiao et al., 2013). Further, the IVS14+3A>T mutation which causes truncation of DIIS2 and S3 of Nav1.1 affects both excitatory and inhibitory neurons (Liu et al., 2013).

In addition to alterations caused by discrete mutations, differences in genetic background represent a major source of variability that complicates interpretation of any experimental findings as they can influence phenotypic differences (Liang and Zhang, 2013; Vitale et al., 2012). Evaluation of a large cohort of control and patient iPSC lines can help assess the contribution of mutations to disease phenotypes (Sandoe and Eggan, 2013). This is problematic when evaluating potential changes in electrophysiological properties that require examination at the single cell level in multiple cells from a large number of cell lines. However, a more direct way to examine the effect of a mutation independent of genetic background is to compare the cellular changes caused by a single mutation in isogenic pairs of control and mutant iPSC lines (Bassett, 2017; Chen et al., 2014; Liu et al., 2016; Sandoe and Eggan, 2013; Smith et al., 2018). A recent study of the DS-associated *SCN1A* Q1923R mutation used transcription activator-like effector nucleases (TALENs) to generate an isogenic corrected line by removing the *SCN1A* mutation (Liu et al., 2016). Comparison of the isogenic corrected versus the patient line revealed a decrease in sodium current density in inhibitory neurons in the patient line. Comparison of the isogenic corrected line with a control line from a different genetic background revealed additional alterations in the voltage dependence of sodium current activation and action potential (AP) threshold suggesting these might be associated with the genetic background of the patient. An isogenic mutant line in which the mutation had been introduced into the control would have been able to distinguish whether this was due to genetic background or interaction between the mutation and the genetic background.

The current study focuses on assessing the contribution of the autosomal dominant *SCN1A* K1270T mutation identified in a GEFS+ family to the etiology of the disease (Abou-Khalil et al., 2001). Two isogenic pairs of iPSC lines generated by CRISPR/Cas 9 gene editing were used: 1a) control (unaffected sibling), 1b) mutated control homozygous for the K1270T mutation, and 2a) patient heterozygous for the K1270T *SCN1A* mutation (GEFS+ sibling), 2b) corrected patient. All 4 iPSC lines in the two isogenic pairs can be differentiated into functional mixed neuronal cultures containing both GABAergic and glutamatergic neurons that expressed Nav1.1. Comparison of electrophysiological properties of neurons within and between pairs identified mutation-dependent alterations independent of genetic background. There were also differences found to be associated with the genetic background and others the result of interactions between the mutation and the background. Such CRISPR/Cas9-based disease modeling strategy with dual isogenic pairs of iPSC-derived neurons represents a promising platform for studying the causality of mutations related to genetic diseases and developing patient-specific therapies based on disease mechanisms.

## 2. Methods

### 2.1 Preparation and maintenance of hiPSCs

Skin fibroblasts were collected from two male siblings of the GEFS+ family by Kevin Ess (Vanderbilt University Medical Center) after obtaining informed consent using IRB approved protocol #080369. The control fibroblast donor had no known clinical diagnoses; the patient sibling had four febrile seizures beginning at 6 months of age and four generalized atypical absence seizures since the age of two (Abou-Khalil et al., 2001). Age at time of biopsy the control sibling was 23 years and the patient sibling was 21 years old.

Skin fibroblasts were reprogrammed into iPSCs using nonintegrating Sendai virus by the Schwartz laboratory as published (Stover et al., 2013). Two human iPSC lines were generated, SC210.12-SF6-2I3.M11S8 (control) and SC215.19-SF4-2I8.M10S6 (patient). iPSCs were maintained in feeder-free conditions following published protocols (Stover et al., 2013; Stover and Schwartz, 2011).

### 2.2 CRISPR/Cas9 editing on iPSCs derived from two siblings

To generate a mutated control line, the K1270T mutation was knocked into an iPSC line generated from the unaffected sibling (SC210.12-SF6-2I3-M11S8-S4-S6-S2). The 20-nt guide RNA (sgRNA) 5’-CTTCTAAAATGGGTGGCATA-3’ was designed using the CRISPR design tool (http://crispr.mit.edu) from the Zhang laboratory and cloned into plasmid pSpCas9(BB)-2A-Puro (Px459) V2.0 (Addgene, 62988). Plasmid constructs were purified with the Endo-free Plasmid Maxiprep kit (Qiagen). The 140-nt repair template ssODN1 5’-GATTAAGACGATGTTGGAATATGCTGACAAGGTTTTCACTTACATTTTCATTCTGGAAATGCTTCTAACATGGGTGGCATATGGATATCAAACATATTTCACCAATGCCTGGTGTTGGCTGGACTTCTTAATTGTTGATG-3’ (Integrated DNA Technologies) was designed with two homologous arms flanking the Cas9 cleavage site, the threonine codon (ACA) at the mutation site and an addition of a silent EcoRV mutation.

To generate a corrected patient line, the mutation was removed from a patient iPSC line (SC215.19-SF4-2I8.M10S6-S2-S2). sgRNA (5’-CTTCTAAC ATGGGTGGCATA-3’) complementary to the target sequence of mutated allele was cloned into the same plasmid. The 140-nt repair template ssODN2 5’-GATTAAGACGATGTTGGAATATGCTGACAAGGTTTTCACTTACATTTTCATTCTGGAAATGCTTCTAAAATGGGTGGCATATGGCTATCAAACATATTTCACCAATGCCTGGTGTTGGCTGGACTTCTTAATTGTTGATG-3’ (Integrated DNA Technologies) was similar to ssODN1 but with the lysine codon at the mutation site (AAA) and without EcoRV silent mutation. 8 × 10^5^ cells were co-transfected with 3 *µ*g plasmid-sgRNA construct and ssODN (0.6 *µ*M final concentration) with Amaxa Human Stem Cell Nucleofector Kit 1 (Lonza) and program A-023 for the Amaxa Nucleofector Device II. To obtain low seeding density, the transfected cell suspension was diluted in a 1:6 – 1:8 ratio and plated into Matrigel (BD Biosciences)-coated wells of 6-well plates. 24 hours post-transfection, cells were treated with 2 *µ*g/ml puromycin (Sigma-Aldrich) in StemPro complete media (Stover and Schwartz, 2011) containing 0.58 *µ*l/ml SMC4 (Corning). 48 hours post-transfection, cells were switched to normal StemPro complete media containing 0.58 *µ*l/ml SMC4. Continued maintenance required media that included 50% conditioned media from 70-90%-confluent unedited iPSC cultures and 50% StemPro complete media with 55 *µ*M β-mercaptoethanol (Invitrogen), 1×StemPro (Invitrogen), 20 ng/ml bFGF (Stemgent) and 0.58 *µ*l/ml SMC4. 6-8 days post-nucleofection, isolated iPSC colonies with 20-30 cells were formed, picked manually using customized glass hooks, and transferred to single wells in 96- or 48-well plates with pre-wet 20 *µ*l barrier micropipette tips. StemPro complete media supplemented with 0.58 *µ*l/ml SMC4 was used for daily feeding. At 100% confluence, clones were split with pre-warmed 62 *µ*l (96-well plate) or 125 *µ*l (48-well plate) of Accutase (Millipore) in a 1:2 ratio. Cells were eventually expanded into two wells in a 48-well plate to be used for freezing as well as sequencing and/or restriction digestion, respectively. All media for feeding and splitting were prepared immediately prior to use and equilibrated at 37°C and 5% CO_2._

A protocol for freezing iPSCs in low density was modified from Stover *et al* (Stover and Schwartz, 2011). Cells at 100% confluence were cryopreserved with pre-chilled freezing medium containing 40% conditioned medium from 70-90% confluent culture, 40% StemPro complete medium, 10% DMSO (Sigma-Aldrich), 10% bovine serum albumin (25% BSA, Gibco) with addition of 0.58 *µ*l/ml SMC4. To increase viability of cells during defrosting, clones from 48-well plates were frozen with 50-250 *µ*l of freezing medium depending on the size of pellets after centrifugation and were stored in 1.2 ml cryo-vials at −80°C.

Genomic DNA was extracted with 10-13 *µ*l of QuickExtract DNA Extraction Solution (Epicentre). Fragments around the intended mutation sites were amplified by PCR in 50 *µ*l reactions with Taq DNA polymerase (Invitrogen), forward primer 5’-CTACCATAGATTCCATCCCCCAA-3’ and reverse primer 5’-TTCCACCAATAGTCTTTCCCCTG-3’. PCR products were confirmed by gel electrophoresis and purified with MinElute PCR Purification Kit (Qiagen) prior to sequencing (Retrogen). MacVector software was used to analyze sequencing data.

Additionally, a subset of mutated control clones was subject to restriction digestion by EcoRV prior to sequencing analysis. As we observed the absence of EcoRV mutation in some of the positive clones with K1270T mutation, direct sequencing was used to screen the remaining clones.

The top five potential off-target sites from CRISPR/Cas9 editing as identified by the CRISPR design tool (http://crispr.mit.edu) from the Zhang laboratory, were *SCN7A*, *FLII*, *SCN2A*, *SCN3A* and *SCN9A*. To determine if any of these were also affected by the editing in the mutated control or the patient corrected line, primers were used to interrogate the relevant region of homology in each gene. PCR conditions were 94 °C, 45 s, 60 °C, 33 s and 72 °C, 55 s for 35 cycles.

To evaluate the expression level of off targets, mRNA was extracted from neuronal cultures of the mutated control line. cDNA was synthesized following the instructions of TRIzol (Thermo Fisher) and SuperScript™ III First-Strand Synthesis SuperMix for qRT-PCR kit (Thermo Fisher), followed by nested PCR. Expression of the off-target gene was compared to *SCN1A* and normalized to *ACTB*. Nested PCR conditions were 94 °C, 45 s, 60 °C, 33 s and 72 °C, 60 s for 35 cycles for all primers. Primers for sequencing and nested PCR are in Supplementary Table 1.

All 4 iPSC lines were karyotyped and cell lines at low passage were expanded for cryopreservation. When samples were thawed for expansion and use in differentiation studies, the *SCN1A* gene was sequenced every 20-30 passages. Research was approved by hSCRO (#2011-1023) and IBC protocols (#2011-1377).

### 2.3 Differentiation of two pairs of isogenic iPSCs into neurons

Two pairs of isogenic iPSCs (4 lines total) within 40 passages of the original generation were patterned into neurons as described (Xie et al., 2018). Neural progenitor cells derived from iPSCs were plated on top of 50-70% confluent mouse astroglial feeder layers that were prepared following a published protocol (Schutte et al., 2018).

At D17 post-plating, excitatory and inhibitory neurons were labeled by transfection with CaMKII-eGFP (Addgene, 50469) and GAD1-mCherry (Genecopoeia, HPRM15652-PM02) plasmids in Opti-MEM (Gibco) and DNA-in (MTI-GlobalStem) or ViaFect (Promega) reagents.

### 2.4 Immunostaining and imaging

To evaluate the expression of pluripotency markers, iPSCs were plated onto glass coverslips coated with Matrigel diluted 1:30 in DMEM/F-12 + Glutamax (Gibco). iPSCs were maintained using StemPro medium (section 2.2). When grown to ∼70% confluence, coverslips were fixed and stained as described (Xie et al., 2018). Primary antibodies OCT-3/4 (1:100; Santa Cruz Biotechnology, sc-5279), SOX2 (1:100; Santa Cruz Biotechnology, sc-365823) and NANOG (1:50; Santa Cruz Biotechnology, sc-293121) were used to characterize the pluripotency of the iPSCs. 4’,6-diamidino-2-phenylindole (DAPI; Life Technologies) staining was added to visualize nuclei.

iPSC-derived neurons were stained with antibodies of βIII-tubulin (1:1000; Sigma-Aldrich, T8660) and GABA (1:4000; Sigma-Aldrich, A2052) to assess the proportion of GABAergic neurons. Neurons were identified by morphology, expression of βIII-tubulin in the cell bodies and neuronal processes, and nuclear localization of DAPI. Identities of neurons labeled by GAD1-mCherry and CaMKII-eGFP plasmids were verified by staining with antibodies of GABA and CaMKII (1:800; Cell Signaling Technology, 3362) respectively. Expression of Nav1.1 was examined in GABA^+^ (1:100; Aldrich-Sigma, A0310) and CaMKII-eGFP^+^ cells by staining with anti-Nav1.1 antibody (1:800; Alomone Labs, ASC-001).

Images were taken by a Zeiss M2 Imager fluorescence microscope and processed using the supplied Zen software.

### 2.5 Whole-cell recording

Whole-cell recordings were obtained from iPSC-derived inhibitory and excitatory neurons at D21-23 post-plating at room temperature. This time period was selected based on studies in two control cell lines demonstrating that a high percentage of neurons were functionally active by this age (Xie et al., 2018). Patch pipettes were pulled from unpolished borosilicate glass (VWR International) by PC-10 pipette puller (Narishige). Open tip resistance was 6-8 MΩ. External solution contained (in mM): 120 NaCl, 5.4 KCl, 0.8 MgCl_2_, 1.8 CaCl_2_, 15 glucose, 20 HEPES, with pH at 7.2-7.4. Osmolality was adjusted to 290 – 295 mOsm. Internal solution contained (in mM): 120 potassium gluconate, 20 NaCl, 0.1 CaCl_2_, 2 MgCl_2_, 1.1 EGTA, 10 HEPES, 4.5 Na_2_ATP with pH at 7.2. Osmolality was around 280 – 283 mOsm. Voltages were corrected for a 5 mV junction potential. To record evoked action potentials, membrane potential was held at −75 mV by injecting sub-threshold currents and cells were depolarized by a series of current injections.

Excitatory and inhibitory postsynaptic currents (EPSCs and IPSCs) were separated at reversal potentials (−49 mV and −2 mV) with a gap-free protocol under voltage clamp. Recordings of spontaneous PSCs (sPSCs) were performed with the standard external solution and a cesium gluconate internal solution containing (in mM): 130 Cs^+^ gluconate, 0.2 EGTA, 2 MgCl_2_, 6 CsCl, 10 HEPES, 2.5 adenosine 5’ triphosphate sodium salt, 0.5 guanosine 5’ triphosphate sodium and 10 disodium phosphocreatine. Voltages were corrected for a 7 mV junction potential. Miniature postsynaptic currents (mPSCs) were recorded in external solution containing 1 *µ*M TTX (tetrodotoxin; Alomones Labs).

Isolated sodium currents were recorded with external solution containing (in mM): 120 NaCl, 5.4 KCl, 0.8 CoCl_2_, 15 glucose, 20 HEPES, 2.5 tetraethylammonium and 1.0 4-aminopyridine. The same cesium gluconate internal solution for PSC recording was used for sodium current recording. Voltages were corrected for a 10 mV junction potential. P/N leak subtraction was turned on. The voltage dependence of activation was determined with a step protocol. Cell membrane potential was increased from a holding potential of −100 mV to a range of potentials between −100 mV and 30 mV for 45 s with a step interval of 5 or 10 mV. Currents recorded at the shared voltage steps were included for current density analysis and G-V graph plotting. Peak transient currents were plotted against holding potentials and fit with the following equation:

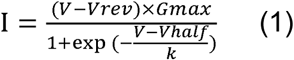

where I is the peak current amplitude, V is the holding potential of each pulse, V_rev_ is the reversal potential, G_max_ is the maximal conductance, V_half_ is the voltage with 50% channels activated, and k is the slope. Current data in 5 mV and 10 mV step intervals were used for fitting. Conductance was calculated with the following equation:

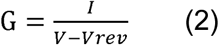

where G is conductance, and I, V, V_rev_ were as described in equation (1).

The voltage dependence of steady-state inactivation was analyzed with a two-step protocol. It began with a holding potential of −100 mV. Next the conditioning pulse was increased through a series of voltage steps from −100 mV to 30 mV for 90 ms with a step interval of 5 or 10 mV. Currents recorded at the shared voltage steps were included for G-V graph plotting. A test pulse at −10 mV for 90 ms was immediately applied and followed by a holding potential of −100 mV. The peak current amplitude during test pulses were normalized to that during the first test pulse and then plotted against the holding potential of the conditioning pulse. The I-V curves were fitted to the following equation:

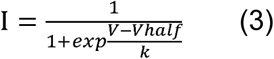

where I is the normalized peak current amplitude during the test pulses, V_half_ is the voltage with 50% channels inactivated, and k is the slope. Current data in 5 mV and 10 mV step intervals were used for fitting.

The recovery from steady-state inactivation was determined using a second two-pulse protocol. Following a holding potential of −100 mV for 27 ms, two test pulses at −10 mV for 90 ms were separated by a time interval with a range of 1-10 ms with an increment of 1 ms in the first protocol, and 10-50 ms with an increment of 5 ms in the subsequent protocol. Peak current amplitude from the second pulse was normalized to the average current amplitude from the first pulse to calculate the fractional recovery. The data of recovery were fitted using the following equation:

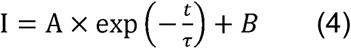

where I is the normalized current amplitude, A is the relative percentage of recovered currents, t is the time interval, and τ is the time constant of recovery.

Electrophysiological data were collected with a List EPC7 amplifier and a Digidata 1320A D-A converter (Axon Instruments), or a MultiClamp 700B amplifier and a Digidata 1440A D-A converter (Axon Instruments). Recordings were obtained by pClamp8 or pClamp10 (Axon Instruments) software with a Bessel filter at 2 kHz and a low-pass filter at 10 kHz.

### 2.6 Data Analysis and Statistics

Counting from fluorescent micrographs was conducted in a blinded fashion with respect to cell line. For each coverslip, at least 3 fields with evenly distributed cells observed via DIC were chosen for counting. 3 coverslips each from individual plating of an iPSC cell line were included for counting the number of cells expressing pluripotency markers.

Electrophysiological data were analyzed using Clampfit 10.6 (Molecular Devices). Cells that were included in electrophysiological analyses had a leak current amplitude smaller than 50 pA at a holding potential of -−75 mV. Evoked action potentials (APs) recorded under current clamp were included if they had a stable baseline at −65 ∼ −85 mV with sub-threshold current injection, a peak amplitude ≥ −5 mV on the first AP and ≥ −15 mV in subsequent APs within the train. AP events were excluded if the maximum firing frequency deviated from mean by more than ± 2SD. Rheobase data were excluded if the resting membrane potentials (RMP) were more depolarized than −35 mV. AP properties were examined in the first AP at rheobase that first elicited AP firing. AP threshold was identified as the first derivative of the membrane potential (dV/dt) 5% or higher than the peak rate of change. AP amplitude was defined as the change of membrane potential from the threshold to the peak. AP amplitude and half-width data were not included if the cell did not fire evoked AP. PSCs recorded under voltage-clamp were analyzed using Mini Analysis 6.0.7 (Synaptosoft). PSCs were identified as events with an amplitude ≥ 5 pA (2× RMS noise of 2.5 pA) and were then manually verified individually. To calculate the (E-I)/(E+I) index, the absolute values of amplitude or charge transfer of EPSCs and IPSCs were used for E and I respectively. Cells were excluded from sodium current analysis if their current density in the range of – 10 – 40mV was 2SD beyond the mean.

Electrophysiological data are presented with n = number of recorded cells and the number of individual platings shown in parentheses. All data were analyzed with D’Agostino-Pearson normality test. Statistical tests were selected based on the normality test results and performed in Prism 7.0.2 (GraphPad Software). Figures were generated using DeltaGraph 7.1 (Red Rock Software).

## 3. Results

### 3.1 Two pairs of isogenic iPSCs generated by CRISPR/Cas9 editing

The autosomal dominant K1270T *SCN1A* mutation identified in a GEFS+ family results in a lysine-to-threonine conversion in the second transmembrane segment of the third homologous domain in the α subunit of the human Nav1.1 protein (Abou-Khalil et al., 2001) (Figure 1A). Skin fibroblasts were collected from two male siblings (an unaffected control and a GEFS+ patient) and both were reprogrammed into iPSC lines using the nonintegrating Sendai virus (Figure 1B). To examine the contribution of the K1270T mutation to disease phenotypes, two pairs of isogenic lines were included in this study. In the first pair, the GEFS+ K1270T mutation (c.4227A>C) was knocked into the unaffected sibling derived control line using CRISPR/Cas9. For the second isogenic pair, the K1270T mutation in the patient line was corrected (c.4227C>A; Figure 1C). The mutated control clone with desired sequence was homozygous, likely the result of the short distance (9 bp) between the Cas9 cut site and the K1270T mutation locus (Paquet et al., 2016).

**Figure 1.**
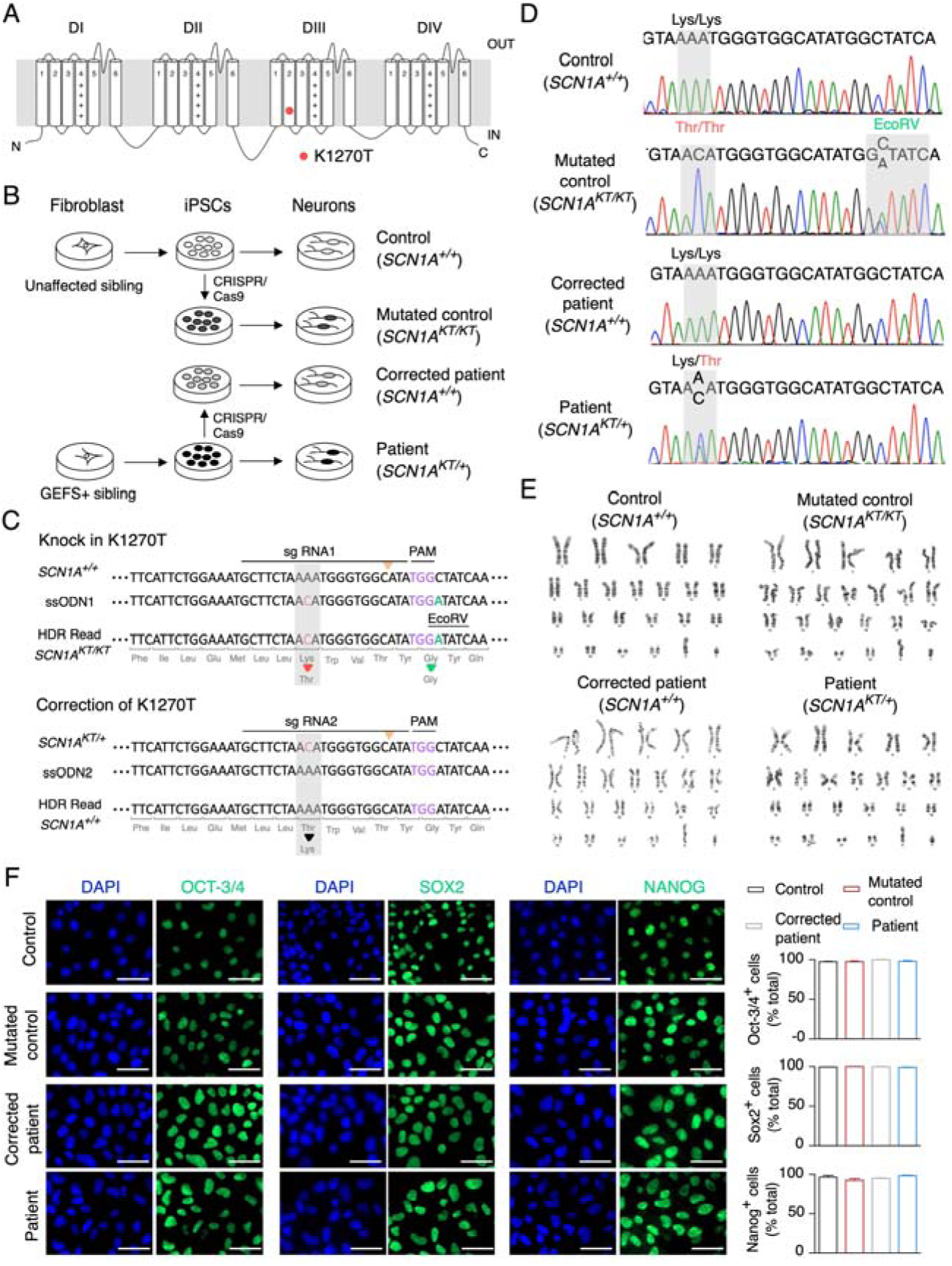
Isogenic pairs of iPSCs generated by CRISPR/Cas9 editing. (A) The K1270T mutation is located in the second transmembrane segment of the third homologous domain of Nav1.1 alpha subunit. (B) CRISPR/Cas9 editing was used to generate two isogenic pairs from iPSCs derived from two siblings in the GEFS+ family (control and patient) followed by differentiation into functional neurons. (C) Scheme of CRISPR/Cas9 editing design. Additional silent mutation introduced by ssODN1 resulted in an EcoRV restriction site for clone screening when knocking in the K1270T mutation. (D) Sequencing of two isogenic iPSC pairs confirmed the absence and the presence of the mutation. (E) All four lines had normal karyotypes (46, XY). (F) iPSCs of two isogenic pairs were stained with nuclei marker DAPI and pluripotency markers OCT-3/4, SOX2 and NANOG individually. The percentage of cells positive for each of the pluripotency markers was not significantly different between the 4 lines (p = 0.1, 0.06 and 0.2 for OCT-3/4, SOX2 and NANOG respectively, one-way ANOVA). Scale bar represents 50 *µ*m. Data represented as mean + s.e.m. Data represent counts from three fields in three coverslips from three platings for each genotype.

Sequencing of DNAs from the iPSC lines in the 2 isogenic pairs confirmed that the control line did not have the mutation at site 1270 (*SCN1A^+/+^*). The mutated control was homozygous for the mutation (*SCN1A^KT/KT^*) based on sequencing and this was further confirmed by the presence of the silent EcoRV mutation from the repair template in only one allele, demonstrating that the editing process did not also result in loss of one allele and therefore, a hemizygous clone. The patient line was confirmed to be heterozygous for the mutation (*SCN1A^KT/+^*) and the mutation was absent in the corrected patient lines (*SCN1A^+/+^*; Figure 1D). The engineered homozygous mutated control provides a unique opportunity to evaluate its cellular defects since two copies of *SCN1A* mutations are usually lethal to organisms *in vivo* (Martin et al., 2010; Yu et al., 2006).

Karyotypes of four iPSC lines were verified normal with 46, XY chromosomes (Figure 1E). Pluripotency of each of the 4 cell lines was confirmed by staining for expression of OCT-3/4, SOX2 and NANOG (Yu et al., 2007) (Figure 1F). The top five potential off-target sites for mutations were evaluated, including *SCN7A*, *FLII*, *SCN2A*, *SCN3A* and *SCN9A* (Supplementary Figure 1A). There were no off-target mutations in candidate genes in the corrected patient line. In the homozygous mutated control line, there was a mutation only in *SCN7A*, in addition to the K1270T knock-in mutation. Since *SCN7A* encodes a non-voltage-gated, TTX-insensitive Na_x_ sodium channel, it is unlikely to affect evaluation of sodium currents and firing properties that are both completely eliminated by addition of TTX to the recording solution (Supplementary Figure 1C). In addition, this gene is expressed in astrocytes, and neurons in circumventricular organs (Gautron et al., 1992; Watanabe et al., 2000) and there is no evidence that this gene is expressed in our iPSC-derived neurons based on nested PCR reactions that resulted in clear amplification of *SCN1A* transcripts with undetectable levels of *SCN7A* (Supplemental Figure 1C).

### 3.2 iPSC-derived neuronal cultures contain both inhibitory and excitatory neurons

We previously optimized a direct differentiation protocol that gives rise to neuronal cultures containing excitatory and inhibitory neurons (Schutte et al., 2018; Xie et al., 2018). With this protocol, iPSCs from both isogenic pairs were differentiated into neural progenitors followed by plating onto astroglial feeder layers for neuronal differentiation. To evaluate the morphology of neurons and quantify the proportion of inhibitory GABAergic and excitatory glutamatergic neurons in the cultures derived from the four iPSC lines, cultures were fixed at D21-24 post plating and stained with anti-βIII-tubulin and anti-GABA or anti-CaMKII antibodies. Cultures from all lines stained βIII-tubulin positive cells with elongated branched neurites (Figure 2A). GABAergic and glutamatergic neurons comprised 30-40% and ∼50% of the total neuronal population respectively, with no significance between genotypes (Figure 2B, p = 0.1, one-way ANOVA for GABAergic neurons and p = 0.6, Kruskal-Wallis test for glutamatergic neurons).

**Figure 2.**
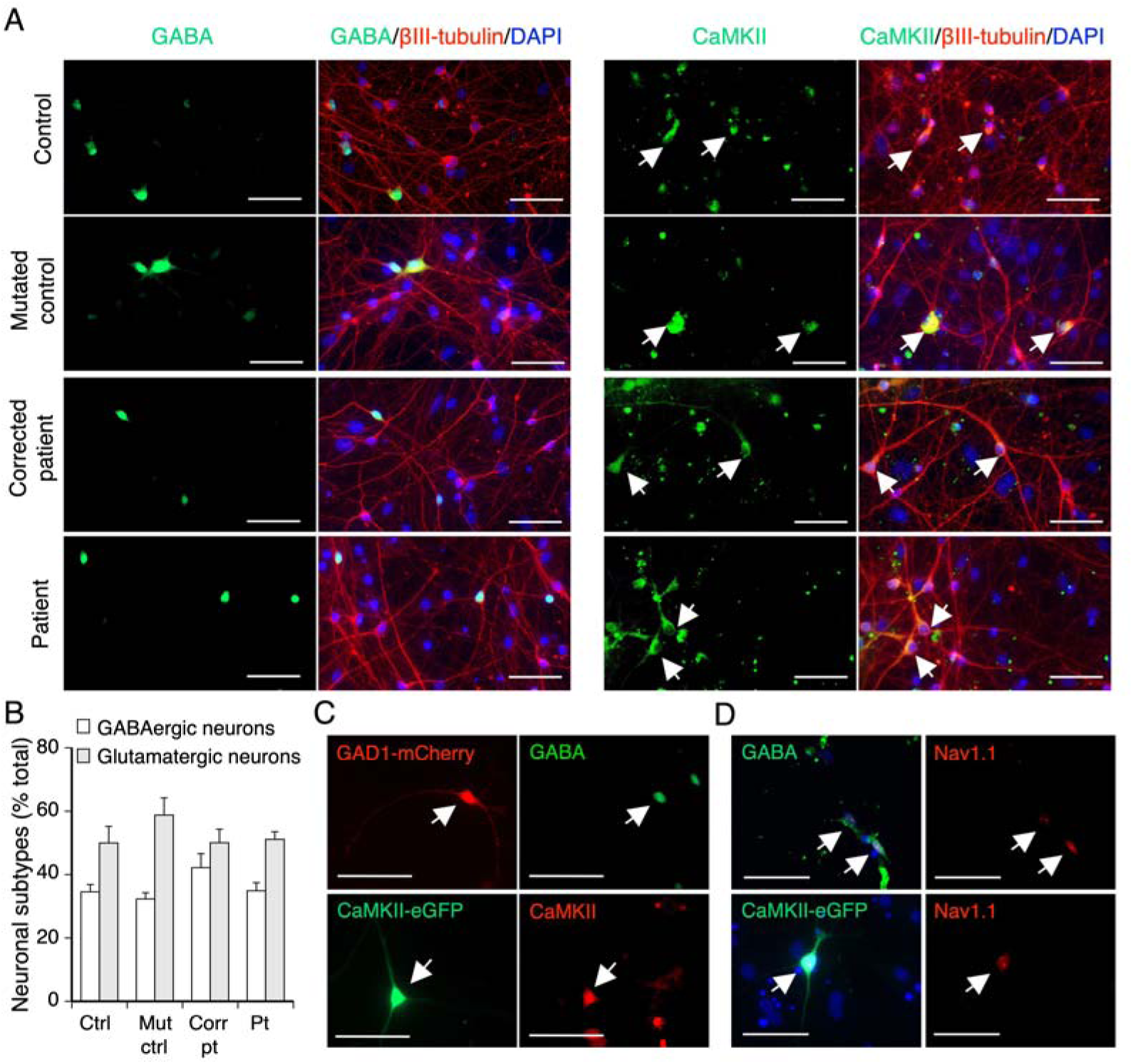
Cultures from two isogenic pairs contain a similar proportion of GABAergic and glutamatergic neurons. (A) Representative micrographs from cultures of two isogenic pairs of iPSC-derived neurons stained with neuronal subtype markers GABA or CaMKII antibodies to identify GABAergic and glutamatergic neurons at D21-24 post plating. (B) The percentage of GABAergic and glutamatergic neurons (white arrows) in neuronal cultures at D21-24 post plating in each line. There was no significant difference between cell lines (p = 0.1, one-way ANOVA for GABAergic neurons and p = 0.6 Kruskal-Wallis test for glutamatergic neurons). (C) Identification of inhibitory (top) and excitatory (bottom) neurons at D21-24 post plating labeled with plasmids via transient transfection during live recording (white arrows). At least 3 coverslips of 2-3 individual platings were evaluated for each line. (D) Localization of Nav1.1 in GABAergic and glutamatergic neurons of the control line (white arrows). Data represented as mean + s.e.m. Ctrl = control, mut ctrl = mutated control, corr pt = corrected patient, and pt = patient. Scale bar represents 50 *µ*m.

In order to identify inhibitory and excitatory neurons during live recording, cells were transiently transfected with GAD1-mCherry and CaMKII-eGFP plasmids at D17 post plating. Visualization of the cultures with fluorescent illumination at D21-24 post plating was used to identify and record from mCherry expressing GAD1-positive inhibitory neurons and eGFP expressing CaMKII-positive excitatory neurons (white arrows, Figure 2C). The plasmid labeling strategy accurately identified GABAergic and glutamatergic neurons based on the colocalization of anti-GABA staining in 90.05% ± 3.26% of GAD1-mCherry^+^ and anti-CaMKII staining in 97.15% ± 2.47% of CaMKII-eGFP^+^ neurons examined. Further, *SCN1A* was expressed in both inhibitory and excitatory neurons of the control and the mutated control lines based on anti-Nav1.1 antibody staining in both GABAergic and glutamatergic neurons (Figure 2D).

### 3.3. Inhibitory neurons: the K1270T mutation reduces evoked firing and sodium current

To determine if the K1270T mutation causes cell type-specific alterations in excitability, evoked action potential recordings were obtained at D21-D24 post plating from iPSC-derived inhibitory neurons in cultures from two isogenic pairs: 1a) control (*SCN1A^+/+^*), 1b) mutated control, homozygous (*SCN1A^KT/KT^*); 2a) corrected patient (*SCN1A^+/+^*), 2b) patient, heterozygous (*SCN1A^KT/+^*). To account for variability between platings, and within plating, data for all electrophysiological properties represent recordings from multiple cells in at least three independent platings, and these values are presented in each figure or table.

Over 90% of inhibitory neurons in each genotype fired action potentials (Figure 3A and 3D). Assessment of firing frequency vs current step curves demonstrated a decrease in mean firing frequency in the homozygous mutated control line compared to the control line (Figure 3B). Both the main effect associated with genotype and an interaction effect for firing frequency were significantly different (genotype: p = 0.0092, interaction: p < 0.0001, two-way ANOVA with repeated measures; Figure 3B and Table 1). The AP amplitude was also significantly smaller in the homozygous mutated control line compared to the control (p = 0.0073, two-tailed unpaired *t* test; Figure 3C).

**Figure 3.**
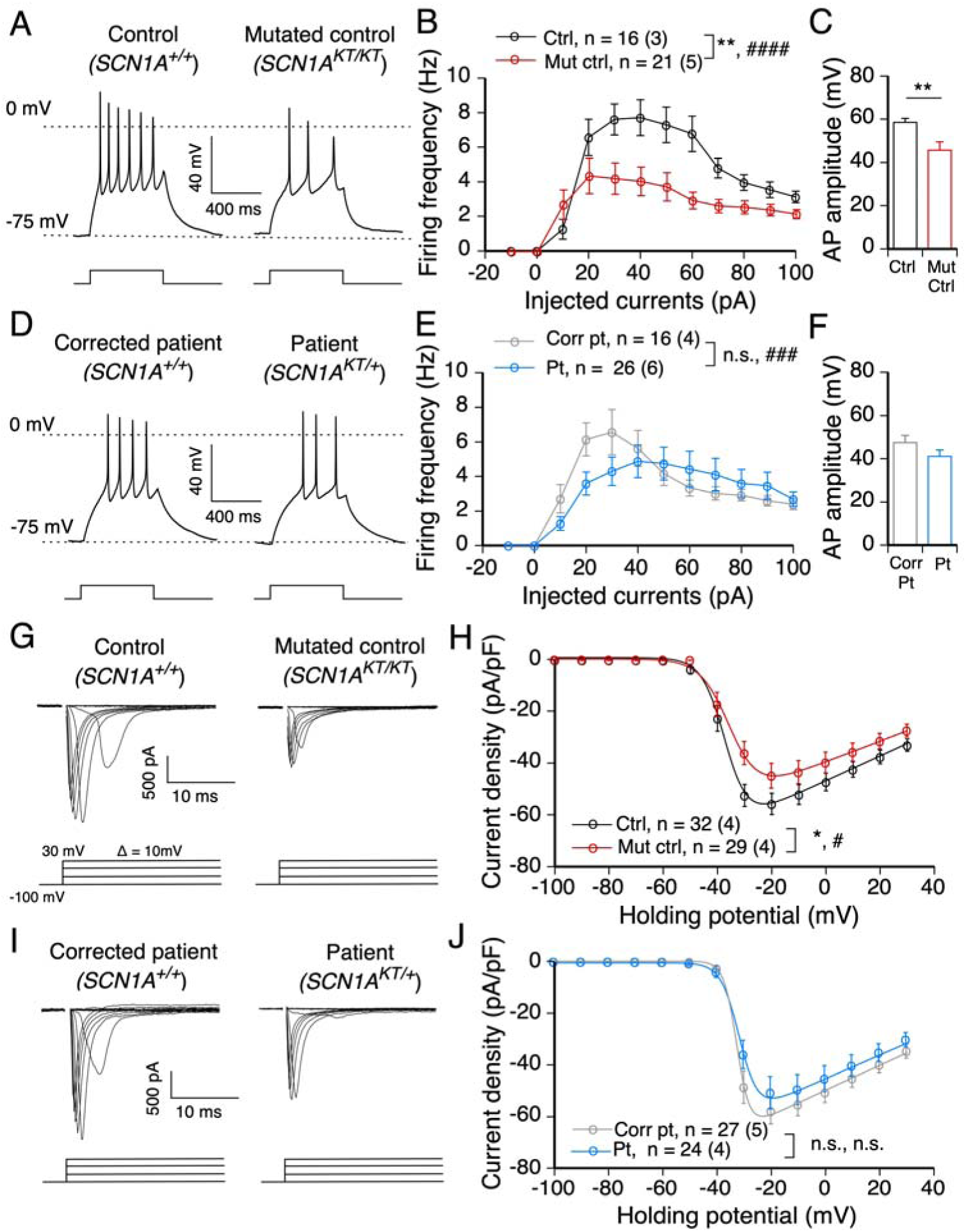
In inhibitory neurons, the K1270T mutation is associated with impaired AP firing and reduced sodium current density. (A) Traces of action potential firing with 30 pA injected in inhibitory neurons of the first isogenic pair of the control (ctrl) and the homozygous mutated control (mut ctrl). (B) The input-output curves of action potentials in inhibitory neurons of the first isogenic pair. (C) AP amplitude of the first AP at rheobase in inhibitory neurons of the first isogenic pair. ** denotes p < 0.001, two-tailed unpaired *t*-test. (D-F) Same as A and B, but in inhibitory neurons of the second isogenic pair of the lines from the corrected patient (corr pt) and the heterozygous patient (pt). (G) Traces of voltage-gated sodium currents in inhibitory neurons of the first isogenic pair. The activation protocol started with a prepulse at −100 mV followed by a series of steps ranging from −100 to 30 mV in 10 mV intervals. (H) Current-voltage relationship in inhibitory neurons of the first isogenic pair. Whole cell currents were normalized to capacitance. (I-J) Same as G and H, but in inhibitory neurons of the second isogenic pair. Data presented as mean + s.e.m in C and F, and mean ± s.e.m with total number of neurons evaluated and the number of individual platings in parentheses for each line in B,E,H, and J. * and # represent significant differences in genotypes and interaction within an isogenic pair respectively, two-way ANOVA and *post hoc* Sidak’s analysis. One to four symbols denote p values < 0.05, 0.01, 0.001, 0.0001 respectively.

**Table 1.** Firing frequency of inhibitory neurons

| Currents<br>(pA) | Control | Mutated<br>control | Mean<br>change<br>(pair 1) | Corrected<br>patient | Patient | Mean<br>change<br>(pair 2) |
| --- | --- | --- | --- | --- | --- | --- |
| -10 | 0 ± 0 | 0 ± 0 | 0 | 0 ± 0 | 0 ± 0 | 0 |
| 0 | 0 ± 0 | 0 ± 0 | 0 | 0 ± 0 | 0 ± 0 | 0 |
| 10 | 1.3 ± 0.6 | 2.7 ± 0.9 | 1.4 | 2.7 ± 0.8 | 1.3 ± 0.4 | 1.4 |
| 20 | 6.6 ± 1.0 | 4.4 ± 1.0 | 2.2 | 6.1 ± 1.0 | 3.6 ± 0.7 | 2.5 |
| 30 | 7.6 ± 0.9 | 4.2 ± 0.9 | 3.4 | 6.6 ± 1.3 | 4.3 ± 0.8 | 2.3 |
| 40 | 7.7 ± 1.0 | 4.0 ± 0.8 | 3.7 | 5.6 ± 1.0 | 4.9 ± 0.9 | 0.7 |
| 50 | 7.4 ± 1.0 | 3.7 ± 0.8 | 3.7 | 4.2 ± 0.7 | 4.7 ± 1.0 | -0.5 |
| 60 | 6.8 ± 1.0 | 2.9 ± 0.5 | 3.9 | 3.3 ± 0.5 | 4.4 ± 1.0 | -1.1 |
| 70 | 4.8 ± 0.6 | 2.6 ± 0.4 | 2.2 | 3.0 ± 0.4 | 4.1 ± 0.9 | -1.1 |
| 80 | 4.0 ± 0.5 | 2.5 ± 0.4 | 1.5 | 2.9 ± 0.3 | 3.6 ± 0.8 | -0.7 |
| 90 | 3.4 ± 0.5 | 2.4 ± 0.3 | 1.0 | 2.6 ± 0.3 | 3.5 ± 0.8 | -0.9 |
| 100 | 3.1 ± 0.3 | 2.1 ± 0.3 | 1.0 | 2.4 ± 0.3 | 2.7 ± 0.4 | -0.3 |

a. Data reported as mean ± s.e.m.
b. Two-way ANOVA and *post hoc* Sidak's analysis

In the heterozygous patient line compared to the corrected patient line, there was no significant decrease in firing frequency associated with the main effect of genotype, but there was a significant interaction effect (genotype: p = 0.067, interaction: p = 0.007, two-way ANOVA with repeated measures; Figure 3E and Table 1). This indicates that the firing frequency differed between the two lines depending on the injected currents. Specifically, the heterozygous patient line had decreased firing frequency at small current steps compared to the corrected patient line. However, at current steps of 50 pA and above, the firing frequency of the patient and the corrected patient line were largely overlapping (Figure 3E and Table 1). The AP amplitude in the heterozygous patient line was smaller than that in the corrected patient line, but the difference was not significant (Figure 3F). The reduction in firing frequency and AP firing amplitude in the heterozygous patient line was approximately half that seen in the homozygous mutated line compared to the control line at the low current steps (Figure 3B and 3E and Table 1). These data suggest that the impairment of action potential firing in inhibitory neurons is gene dose-dependent. There were no differences in AP threshold, rheobase, the injected current that first elicited AP firing, and half width within and between isogenic pairs (Table 2).

**Table 2.**
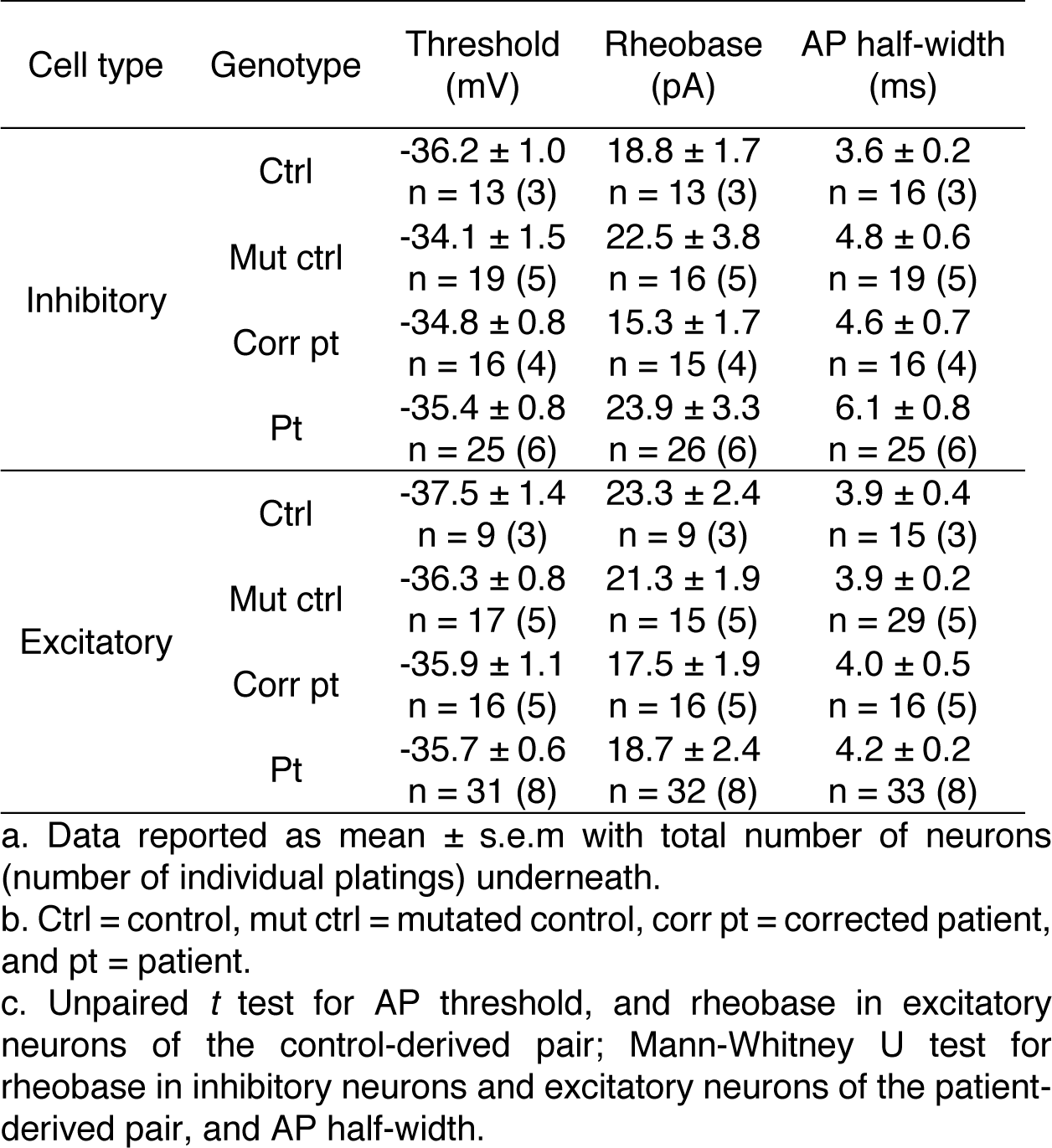
Measurements of excitability in two isogenic pairs

The control and the mutated control had similar input resistance (IR), whole cell capacitance (C_m_) and rest membrane potential (RMP; Table 3). Interestingly while the patient line had comparable input resistance to the isogenic corrected line, it had a significantly smaller whole cell capacitance and a more depolarized RMP (p = 0.024, and < 0.0001 respectively, Mann-Whitney U test for C_m_ and unpaired *t* test for RMP; Table 3).

**Table 3.**
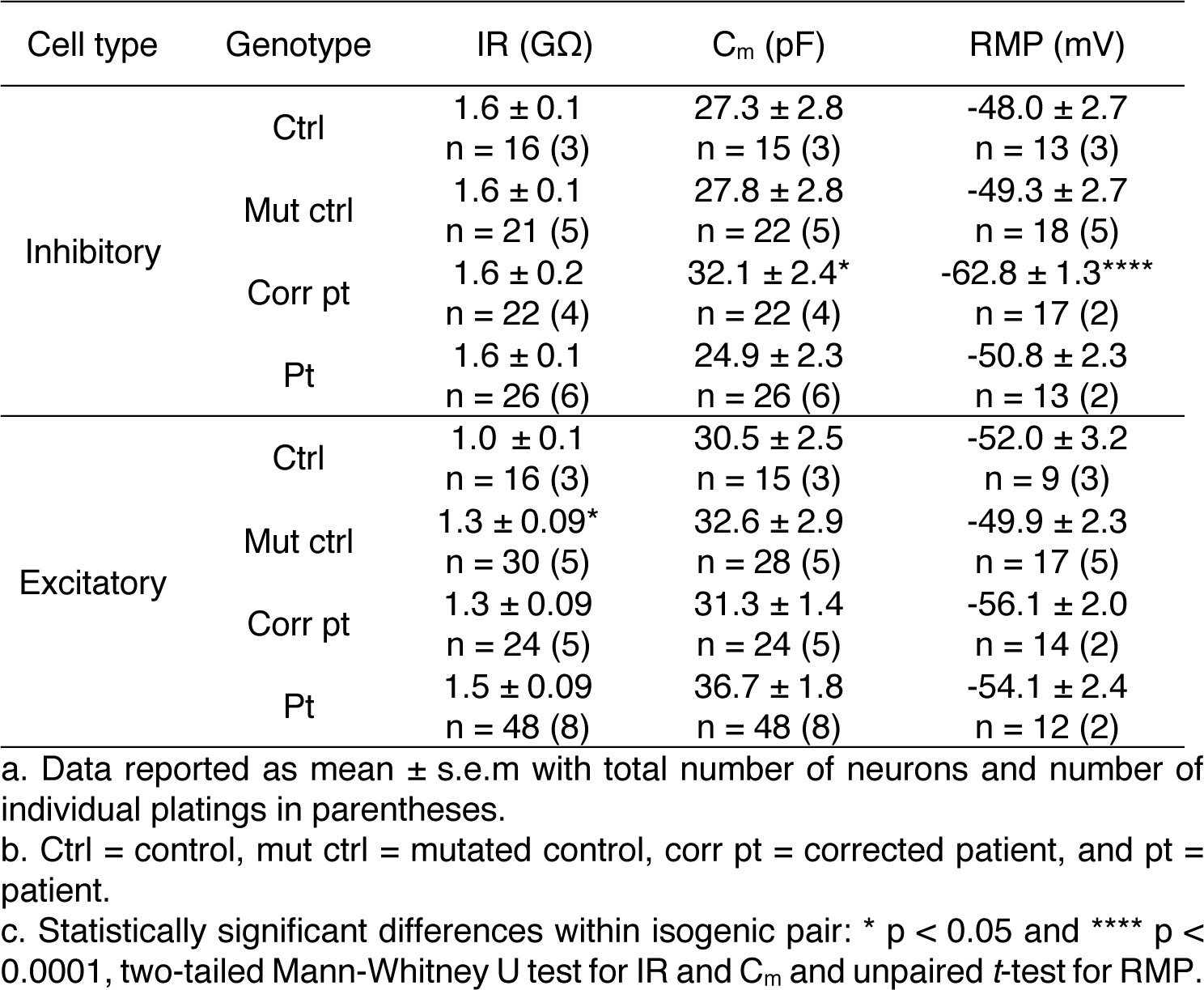
Comparison of passive membrane properties in two isogenic pairs

To examine whether reduced firing frequency and AP amplitude is associated with alterations in sodium current, isolated sodium currents were recorded in inhibitory neurons at D21-24 post plating. All inhibitory neurons in each genotype expressed sodium currents. Sodium currents were not well space clamped at this age in culture in any of the lines (Figure 3G and 3I), consistent with the localization of some sodium channels at sites electrically distant from the neuronal cell bodies as is typical of neurons that fire multiple APs. This is also consistent with many other studies in which iPSC-derived neurons that are capable of firing a train of APs do not have well clamped sodium currents (Jiao et al., 2013; Kim et al., 2018; Liu et al., 2016, 2013; Sun et al., 2016). However, comparative analysis of sodium current properties between control and patient lines in these studies, as in other published studies, provide insights into the disease mechanism. Therefore, our study focuses on comparing sodium current properties from a family of traces within and between isogenic pairs. While the absolute values may not be reflective of the biophysical properties of sodium currents measured from single channels or in expression systems where there is good voltage control, they allow identification of properties that are or are not associated with the mutation in the native environment.

The mean sodium current density versus voltage curves, generated from analysis of inhibitory neurons in each line of an isogenic pair, were compared. The sodium current density in the homozygous mutated control line was reduced compared to the control line. Both the main effect of genotype and an interaction were significant (genotype: p = 0.050, interaction: p = 0.016, two-way ANOVA with repeated measures; Figure 3G and 3H). In the patient line, heterozygous for the K1270T mutation, there was a similar trend towards a decrease in mean sodium current density at each voltage step above −30 mV compared to the corrected line but the main effect associated with the genotype and the interaction effect were not significantly different (genotype: p = 0.27, interaction: p = 0.33, two-way ANOVA with repeated measures; Figure 3I and 3J).

No differences were observed in the voltage-dependence of activation, steady-state inactivation, or recovery from inactivation within the isogenic pairs (the mutated control line compared to the control; the patient compared to the corrected patient line; Figure S2). However, comparison between the isogenic pairs revealed a ∼4 mV depolarized shift in voltage dependence of activation in the pair derived from the control compared to the pair derived from the patient (genotype: p = 0.0065, interaction: p < 0.0001, two-way ANOVA with repeated measures; Table 4 and Figure S2A and S2B). There was also a 3 mV depolarized shift in the voltage dependence of steady-state inactivation in the control vs the patient derived lines (genotype: p = 0.28, interaction: p < 0.0001, two-way ANOVA with repeated measures; Table 4 and Figure S2C and S2D). Shifts between pairs are likely associated with the genetic background of the GEFS+ patient which would not be found if using unrelated control and patient cell lines. Time constant of recovery from inactivation was similar within isogenic pairs (Table 4 and Figure S2E and S2F).

**Table 4.** Comparison of Voltage dependence of Nav in isogenic pairs

| Cell type | Line | Activation |  | Inactivation |  | Recovery |
| --- | --- | --- | --- | --- | --- | --- |
|  |  | V<sub>1/2</sub> (mV) | k (mV) | V<sub>1/2</sub> (mV) | k (mV) | τ (ms) |
| Inhibitory | Ctrl, n = 32 (4) | -38.2 ± 1.4 | 1.7 ± 0.3 | -44.0 ± 1.0 | 5.5 ± 0.2 | 3.0 ± 0.2 |
|  | Mut ctrl, n = 29 (4) | -35.1 ± 1.4 | 2.1 ± 0.3 | -43.3 ± 0.6 | 6.3 ± 0.3 | 3.2 ± 0.2 |
|  | Corr pt, n = 27 (5) | -32.0 ± 0.9 | 2.2 ± 0.3 | -41.0 ± 0.4 | 6.3 ± 0.3 | 2.8 ± 0.1 |
|  | Pt, n = 24 (4) | -31.3 ± 1.3 | 2.5 ± 0.4 | -41.0 ± 0.8 | 6.1 ± 0.3 | 2.9 ± 0.2 |
| Excitatory | Ctrl, n = 36 (4) | -35.4 ± 1.1 | 1.7 ± 0.2 | -43.2 ± 0.5 | 5.7 ± 0.1 | 3.0 ± 0.1 |
|  | Mut ctrl, n = 27 (4) | -35.3 ± 1.3 | 2.2 ± 0.3 | -44.0 ± 0.9 | 5.8 ± 0.2 | 3.4 ± 0.2 |
|  | Corr pt, n = 26 (5) | -31.5 ± 1.0* | 2.3 ± 0.3 | -42.5 ± 0.7* | 7.2 ± 0.5** | 2.9 ± 0.1 |
|  | Pt, n = 26 (4) | -34.7 ± 1.0 | 2.0 ± 0.3 | -40.5 ± 0.4 | 5.4 ± 0.2 | 3.1 ± 0.1 |

a. Data reported as mean ± s.e.m with total number of neurons and number of individual platings in parentheses for each line.
b. Ctrl = control, mut ctrl = mutated control, corr pt = corrected patient, and pt = patient.
c. Statistically significant differences within isogenic pairs: \* p < 0.05 and \*\*\* p < 0.001, two-tailed unpaired t-test for activation and inactivation V<sub>1/2</sub>, and Mann-Whitney U test for inactivation k.

Overall the differences within the isogenic pairs demonstrate that the K1270T mutation causes reduction in evoked firing, action potential amplitude and sodium current density recorded at the cell body in inhibitory neurons. The alterations observed in voltage dependence of activation and steady-state inactivation of sodium currents between isogenic pairs suggest these changes are associated with the genetic background, independent of the mutation.

### 3.4 Excitatory neurons: the K1270T SCN1A mutation reduces sodium current density but does not reduce firing

Previous studies in iPSC and mouse models have suggested that *SCN1A* mutations reduce evoked firing and sodium currents specifically in inhibitory neurons (Hedrich et al., 2014; Martin et al., 2010; Sun et al., 2016; Yu et al., 2006). However, some studies in other *SCN1A* mutations showed excitability and sodium currents of both excitatory and inhibitory neurons are affected by *SCN1A* mutations (Jiao et al., 2013; Liu et al., 2013; Mistry et al., 2014). Antibody staining demonstrated that both excitatory and inhibitory iPSC-derived neurons in our cultures express Nav1.1 channels. Therefore, it was important to evaluate evoked firing and sodium currents in iPSC-derived excitatory neurons in the same two isogenic pairs examined in the previous section at D21-24 post plating.

Over 93% of excitatory neurons in each genotype fired action potentials with injection of depolarizing currents (Figure 4A and 4D). There was no significant main effect difference or interaction effect in the firing frequency versus injected current curves in the homozygous mutated control compared to the control (genotype: p = 0.91, interaction: p = 0.57, two-way ANOVA with repeated measures; Figure 4B). However, in the heterozygous patient line the firing frequency was significantly increased compared to the corrected patient line at current injections over 30 pA (genotype: p = 0.041, interaction: p = 0.0042, two-way ANOVA with repeated measures; Figure 4E). Since the increased firing frequency in excitatory neurons is only in the patient-derived line and not in the isogenic corrected line or in the control-derived pair (control and mutated control lines), this suggests the phenotype is associated with an interaction between the mutation and the patient genetic background. There was no significant difference in the AP amplitude, threshold, rheobase and AP half-width within or between the isogenic pairs (Figure 4C, 4F and Table 2).

**Figure 4.**
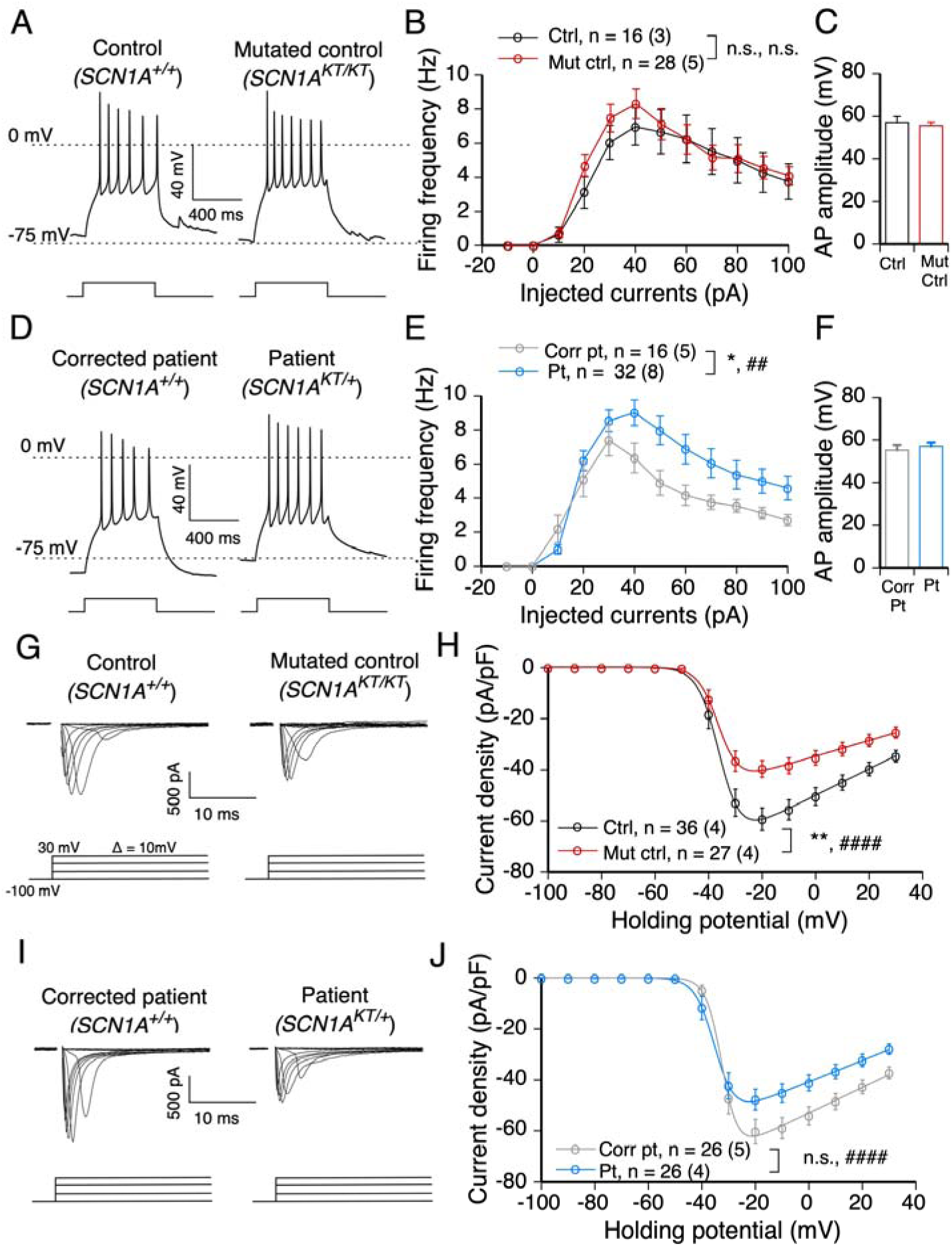
In excitatory neurons, the K1270T mutation affects AP firing and decreases sodium current density. (A) Traces of action potential firing with 30 pA injected in excitatory neurons of the first isogenic pair of the control (ctrl) and the homozygous mutated control (mut ctrl). (B) The input-output curves of action potentials in excitatory neurons of the first isogenic pair. (C) AP amplitude of the first AP at rheobase in excitatory neurons of the first isogenic pair. (D-F) Same as A and B, but in excitatory neurons of the second isogenic pair of the lines of corrected patient (corr pt) and the heterozygous patient (pt). (G) Traces of voltage-gated sodium currents in excitatory neurons of the first isogenic pair. The activation protocol started with a prepulse at −100 mV followed by a series of steps ranging from −100 to 30 mV with a 10 mV interval. (H) Current-voltage relationship in excitatory neurons of the first isogenic pair. Whole cell currents were normalized to capacitance. (I-J) Same as G and H, but in excitatory neurons of the second isogenic pair. Data presented as mean + s.e.m in C and F, and mean ± s.e.m with total number of neurons evaluated and the number of individual platings in parentheses for each line in B,E,H, and J. * and # represent significant differences in genotypes and interaction within an isogenic pair respectively, two-way ANOVA and *post hoc* Sidak’s analysis. One to four symbols denote p values < 0.05, 0.01, 0.001, 0.0001 respectively.

The input resistance in the mutated control was slightly higher than that of the control (p = 0.025, two-tailed Mann-Whitney U test; Table 3). However, there was no difference in the input resistance between the patient vs the corrected patient line (Table 3). The whole cell capacitance and RMP were similar within the isogenic pairs (Table 3).

Based on the lack of decrease in firing in excitatory neurons, and the increase in firing in the patient line, it seemed likely there would be no decrease in sodium current density in the mutated control and possibly an increase in the patient line. To evaluate this, isolated sodium currents were recorded at D21-24 post plating and like inhibitory neurons, all excitatory cells expressed sodium currents that were not well clamped (Figure 4G and 4I). Unexpectedly the current density vs voltage curve demonstrated a significant reduction in sodium current density recorded in the cell body in the homozygous mutated control line compared to the control line (genotype: p = 0.0025, interaction: p < 0.0001, two-way ANOVA with repeated measures; Figure 4H). The sodium current density in the heterozygous patient line was also smaller compared to the corrected patient line. While the main effect of genotype was not significant, there was a significant interaction effect indicating that sodium current density differed between the two lines depending on the voltage steps (genotype: p=0.056, interaction: p < 0.0001, genotype, two-way ANOVA with repeated measures; Figure 4J). The reduction of sodium current density in the heterozygous patient line at the higher voltage steps was approximately half the magnitude seen in the homozygous mutated control line (Figure 4H and 4J). This is consistent with a gene dose-dependent effect of the mutation.

Since there was no concomitant reduction in firing frequency in the excitatory neurons in the homozygous mutant line, this suggests that the properties of other channels, for example potassium channels, in excitatory neurons are more influential in determining the firing phenotype. However, this does not address how a decrease in sodium current in the patient line is consistent with a small but significant increase in excitatory neuron firing frequency. Comparison of other sodium current properties revealed two additional changes in the heterozygous patient line that were not seen in any of the other three lines. There was a hyperpolarized shift in the voltage-dependence of activation and a depolarized shift in steady-state inactivation in the heterozygous patient compared to the corrected patient line (p = 0.028 and 0.015 respectively, two-tailed unpaired *t* test; Table 4 and Figure S3B and S3D). These alterations are both likely to lead to increased channel opening probability over a larger voltage range that would result in an increased evoked firing rate only in this line (Figure 3F). The differences in activation and steady-state inactivation in only one of the four lines indicate that it is due to an interaction between the K1270T mutation and the patient genetic background.

Taken together, similar to the inhibitory neurons, the K1270T mutation causes reduced sodium current density in excitatory neurons. However, the decreased sodium current density caused by the mutation in excitatory neurons is not sufficient to reduce the evoked firing or AP waveforms in the mutated control line. Further, interactions between the mutation and the patient background appear to alter other properties of the sodium channel that increase firing in excitatory neurons of the patient line which may contribute to increased network activity.

### 3.5. The K1270T mutation results in a hyperexcitable network

A previous study from our laboratory found that the large majority of neurons in iPSC-derived cultures receive spontaneous glutamatergic and GABAergic synaptic input at D21-24 post plating (Xie et al., 2018). To determine if the K1270T mutation, that specifically reduces firing of inhibitory neurons or increases firing of excitatory neurons in the patient line, causes an increase in excitability of the neuronal network, spontaneous excitatory and inhibitory post-synaptic currents (sEPSCs and sIPSCs) were recorded from excitatory neurons in each genotype.

All neurons that received both sEPSCs and sIPSCs were included for analysis. Recordings were obtained at two different holding potentials, −2 mV and −49 mV in each neuron corresponding to the reversal potential for EPSCs and IPSCs, respectively (Figure 5A and 5D). To determine whether there was any change in the balance of excitation vs inhibition, an index, (E-I)/(E+I) was used, where E and I represented amplitude or charge transfer of sEPSC and sIPSC respectively in excitatory neurons. This symmetric index ensures that E and I have the same scale, in contrast to an E/I ratio where the range is asymmetrical. An increase in the (E-I)/(E+I) index indicates an increase in excitatory drive or a decrease in inhibitory drive, and vice versa.

**Figure 5.**
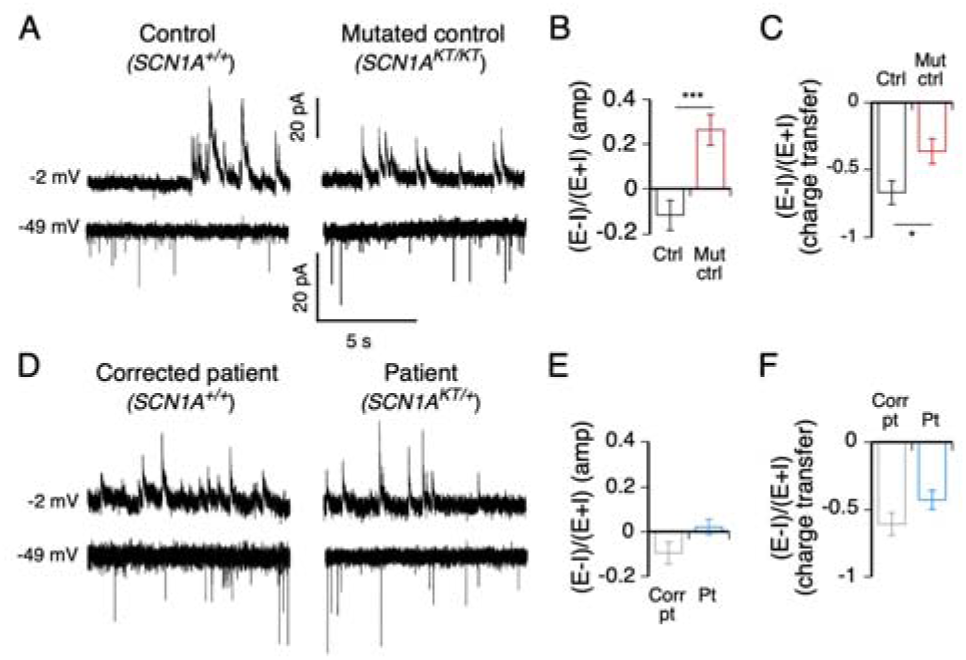
The K1270T mutation gives rise to a hyperactive neural network. (A) Spontaneous excitatory and inhibitory post-synaptic currents received by excitatory neurons in the first isogenic pair of the control (ctrl) and the mutated control (mut ctrl) line. (B) The ratio of spontaneous (E-I)/(E+I) amplitude in excitatory neurons of the first isogenic pair. (C) The ratio of spontaneous (E-I)/(E+I) of charge transfer in excitatory neurons of the first isogenic pair. (D-F) Same as A-C, but in excitatory neurons of the second isogenic pair of the corrected patient (corr pt) and the patient (pt) lines. Data reported as mean ± s.e.m. The total number of neurons and number of platings in parentheses for all four lines are 29 (7), 17 (4), 13 (3), and 18 (3) respectively. *p < 0.05 and ***p < 0.0001, two-tailed Mann-Whitney U test.

The (E-I)/(E+I) index based both on synaptic current amplitude and charge transfer, was significantly larger in the homozygous mutated control line compared to the control and these differences were significant (p = 0.0002 and 0.046 respectively, two-tailed Mann-Whitney U test; Figure 5B and 5C). There was a similar trend of larger (E-I)/(E+I) index in the heterozygous patient line compared to the corrected patient line but these differences were not significant (Figure 5E and 5F). This indicates the mutation results in increased excitability in the neural networks formed in these cultures, consistent with the effects of mutation on sodium currents and evoked firing in inhibitory neurons. It also suggests that the increased firing frequency in the excitatory neurons in the patient line does not significantly affect network activity.

To test whether the increased excitatory drive in the network was also influenced by alterations in synaptic vesicle release, miniature excitatory and inhibitory PSCs (mEPSCs and mIPSCs) were recorded in excitatory neurons in the presence of 1 *µ*M tetrodotoxin to block action potentials. The frequency, amplitude, and charge transfer of mEPSCs were similar between genotypes with the same genetic background (Figure S4A-C). There were also no changes in the mIPSCs (Figure S4D-F). This suggests that the K1270T mutation does not affect the synaptic vesicle release.

Together these data indicate that the K1270T mutation results in an imbalance in excitation and inhibition that leads to hyperactivity in the neural network.

## 4. Discussion

### 4.1 Advantages of dual isogenic pair approach

Previous studies have used pairs of isogenic human iPSCs to explore relationship between gene mutations and phenotypes (Ananiev et al., 2011; Bellin et al., 2013; Smith et al., 2018). To our knowledge, ours is the first study to use two pairs of isogenic iPSCs (control sibling vs mutated control, and corrected patient vs patient sibling) to examine the direct relationship, independent of genetic background, between a single ion channel mutation and cellular mechanisms of genetic epilepsy. This approach is an efficient strategy to evaluate the causality of single gene mutations that require analysis at the single cell level to understand cellular mechanisms as is the case for the many different ion channel mutations linked to epilepsy.

Mutations that are not only necessary but sufficient to cause the functional change are identified as alterations that occur in both mutant lines, the patient line that carries the endogenous mutation and the mutated control line where K1270T was introduced by CRISPR, and not in the control or corrected patient lines that both lack the mutation but the corrected line has been subjected to CRISPR. In the present study, decreases in firing frequency in inhibitory neurons, and sodium currents in excitatory neurons, in both mutant lines and not in the isogenic controls indicate these phenotypes are due to the K1270T mutation, independent of genetic background. Other changes, AP amplitude and sodium current density in inhibitory neurons and excitatory synaptic drive in excitatory neurons were significant only in the homozygous mutant line compared to its isogenic control. However, changes in the same direction for these properties that were smaller magnitude in the heterozygous line compared to its isogenic control suggest a dose-dependent effect of the K1270T mutation (Table 5). A limitation of this interpretation is that the homozygous and heterozygous mutations are in different genetic backgrounds. To determine how the zygosity of the mutation affects the neuronal activity in the same genetic background, future studies will include generation of a heterozygous mutated control line and a homozygous patient line using CRISPR/Cas9 editing to compare with the existing lines.

**Table 5.**
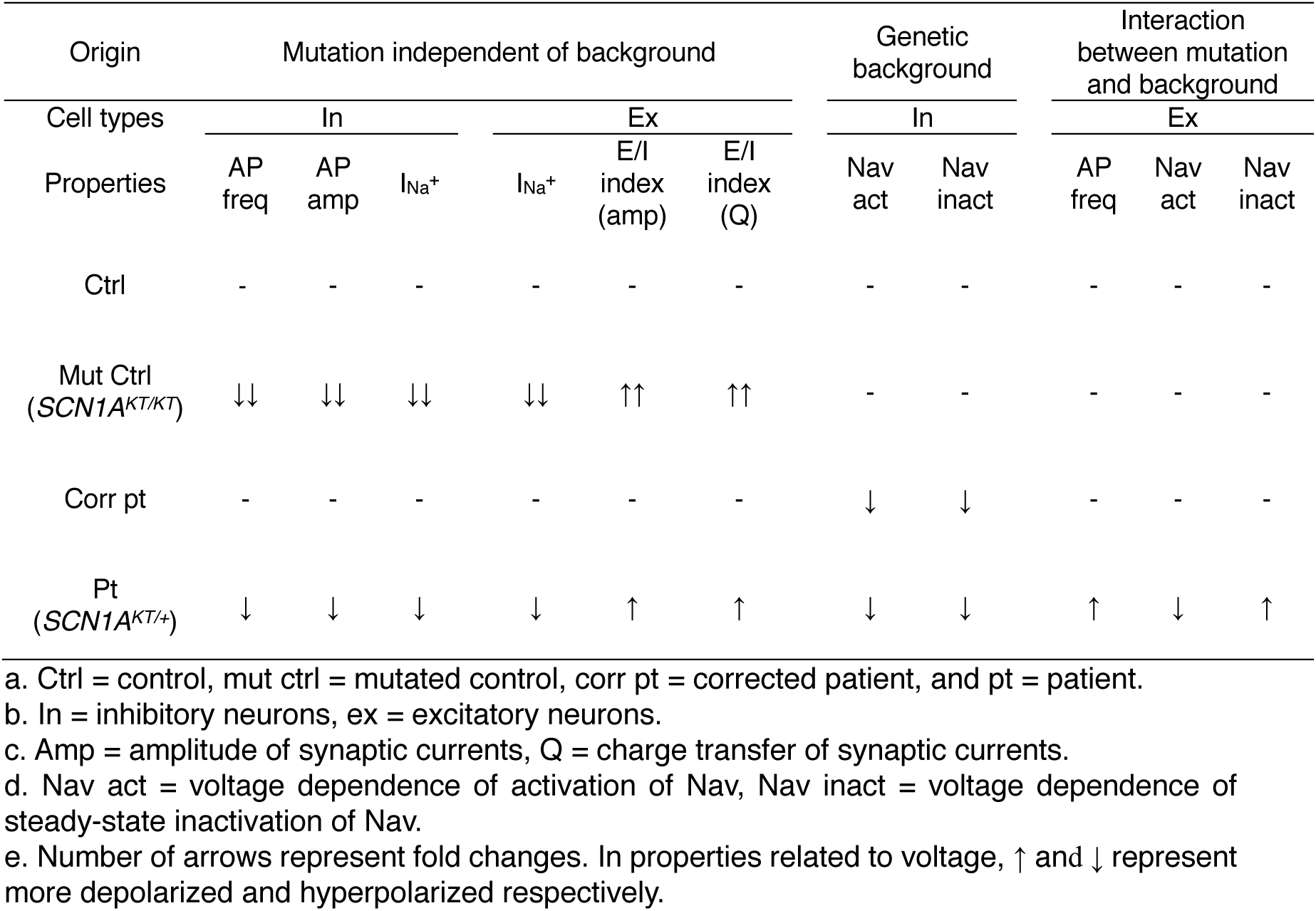
Origins of effects

This comparative strategy also allows for identification of alterations due to the genetic background based on changes that are found in both lines of one isogenic pair but not in the other isogenic pair (Smith et al., 2018), as is seen in the voltage dependence of Nav activation and inactivation in inhibitory neurons of the patient-derived pair only (Table 5). Further, a change in one mutant line that is not seen in the other mutant line or the controls, identifies alterations associated with an interaction between the mutation and the genetic background, as is observed in the voltage dependence of Nav activation and inactivation and increased firing frequency in excitatory neurons of the patient line (Table 5).

The GEFS+ patient from whom we collected skin fibroblast had prolonged febrile seizures and generalized afebrile absence seizures that are more severe than other members with the same *SCN1A* mutation in the pedigree (Abou-Khalil et al., 2001). It is likely that genetic background-dependent functional alterations that were observed in the patient line in the current study contribute to the differences in seizure phenotypes between patients in the GEFS+ family. It will be interesting to compare the genetic background of this individual to other GEFS+ family members to identify specific genetic modifiers that exacerbate or ameliorate the cellular defects. For instance, a missense mutation in *SCN9A* that encodes Nav1.7 has been identified in other *SCN1A*-associated epilepsy patients (Singh et al., 2009). To identify genetic modifiers that are altered in the *SCN1A* GEFS+ mutants, single cell RNA sequencing or exome sequencing will be useful for screening candidates in a high-throughput manner (Hawkins et al., 2016; Landis et al., 2017).

Transcriptomic analysis in multiple iPSC lines generated from different individuals reveals genetic background variation accounts for differences between lines (Burrows et al., 2016; Rouhani et al., 2014). Therefore, it will be interesting to generate additional cell lines and multiple clones from each line via genome editing to robustly interrogate the interaction between the mutation and genetic background in determining cellular phenotypes.

### 4.2 Alterations in excitability and sodium current density in inhibitory neurons

To examine the cell type-specific effect of the mutation in the electrophysiological properties, inhibitory and excitatory neurons are labeled by transfection of the GAD1-mCherry and CaMKII-eGFP plasmids respectively. Specificity of the labeling was verified by immunostaining with their own markers (Figure 2C). It is unlikely that the majority of the iPSC-derived neurons are of an intermediate phenotype, expressing both GABA and CAMKII, for the following reasons. (1) GABAergic and glutamatergic neurons account for ∼35% and ∼55% of total neuronal population. If there was a large percentage of neurons that co-express both markers, we would expect to see their additive percentages exceed 100%. (2) The proportion of different neuronal subtypes in cultures is consistent with data from the pharmacology experiment in our previous study in which we found the majority of synaptic connections in neural networks are GABAergic and glutamatergic, and a small population is glycinergic (Xie et al., 2018). (3) Exposure of cultures to a mixture of anti-GABA and anti-CaMKII antibodies resulted in co-labeling with both antibodies in 17.8% of GABA+ neurons and 6.7% of CaMKII+ neurons. These were unlikely to be represented in the neuronal population targeted for recording based on their size being less than half of the minimum size targeted for whole cell recordings. Together these data indicate the plasmids label neuronal subtypes with high specificity.

Our results demonstrate that in inhibitory neurons, the K1270T *SCN1A* mutation causes decrease in evoked firing frequency and AP amplitude. The reduced excitability and sodium currents in inhibitory neurons is consistent with a number of studies in mouse and iPSC model systems (Martin et al., 2010; Sun et al., 2016; Yu et al., 2006). In particular, previous studies have shown impaired excitability in two subtypes of GABAergic neurons, parvalbumin (PV)- and somatostatin (SST)-expressing interneurons in mouse models of *SCN1A* epilepsy (Favero et al., 2018; Tai et al., 2014). Selective deletion of *Scn1a* in PV and SST interneurons causes hyperactivity in neural networks and seizure behaviors, suggesting the importance of Nav1.1 in regulating inhibitory neuron activity and the E/I balance (Cheah et al., 2012; Dutton et al., 2013; Rubinstein et al., 2015). Our preliminary data reveal that GABAergic neurons in our cultures can be divided into at least two groups staining iPSC-derived neurons with anti-βIII-tubulin and PV or SST antibodies (data not shown). However, we did not observe fast spiking, bursting or adapting firing patterns characteristic of PV and SST interneurons in recorded GAD1-mCherry^+^ neurons. This suggests that functional maturation of interneuron subtypes require a longer time in culture. Future studies will be directed to determine the effect of *SCN1A* mutation in the electrophysiological properties of identified PV and SST interneurons.

iPSC-derived inhibitory neurons in the present study had large sodium currents, but they were not well space clamped. This is typical in neurons that are capable of firing multiple APs (Jiao et al., 2013; Kim et al., 2018; Liu et al., 2016, 2013; Sun et al., 2016). While evaluation of sodium currents that are not well space clamped does not resolve absolute biophysical properties, comparisons of sodium current properties within and between isogenic pairs are still informative in identifying alterations caused by the mutation. For example, pair wise comparisons demonstrate that the K1270T mutation causes a decrease in sodium current density in inhibitory neurons. Further studies will be necessary to determine if the reduction in sodium current density recorded at the cell bodies is due to a decrease in total number of sodium channels expressed or a change in localization of channels as channels distant from the cell soma might reduce their contribution to the recorded currents. This could be addressed by immune-electron microscopy to evaluate voltage-gated sodium channel localization in neurons from the different genotypes.

### 4.3 Alterations in excitability and sodium current density in excitatory neurons

Previous studies have reported mutation of *SCN1A* has little effect on sodium currents in excitatory neurons (Martin et al., 2010; Sun et al., 2016; Yu et al., 2006). In contrast we report that the K1270T mutation results in a similar decrease in sodium current density in excitatory and inhibitory neurons. This is consistent with Nav1.1 staining not only in GABAergic neurons but in our CaMKII-eGFP^+^ neurons. While previous studies suggest that Nav1.1 is strongly expressed in inhibitory neurons (Ogiwara et al., 2007), other studies report that Nav1.1 is present in a subset of excitatory neurons (Dutton et al., 2013; Ogiwara et al., 2013).

Interestingly, the decrease in sodium current density does not result in reduced firing in the excitatory neurons as it does in inhibitory neurons. This indicates that the changes in the sodium current density were not sufficient to reduce excitability in this cell type. This could be due to compensation from potassium currents, as a voltage-gated potassium channel Kv8.2 has been shown to function as a genetic modifier increasing the susceptibility of an *Scn2a* mutant mouse to epilepsy (Bergren et al., 2009; Jorge et al., 2011). Altered currents of Kv1 and Kv4 are associated with increased excitability in a subset of pyramidal neurons in a mouse model of fragile X syndrome (Kalmbach et al., 2015). *CACNB4* and *CACNA1A* that encode Cav have also been identified as genetic modifiers that result in a gain of function in families with *SCN1A* Dravet syndrome (Ohmori et al., 2013, 2008).

Further, the AP firing frequency is actually increased in the excitatory neurons in the patient line resulting from an interaction between the mutation and the patient genetic background. This is consistent with changes in the voltage-dependence of sodium current activation and inactivation only in the patient excitatory neurons. These changes predict that the channel is open over a wider voltage range and is thus consistent with the increase in AP firing seen in patient excitatory neurons. To explore the biophysical changes in finer detail it would be possible to lower the external sodium concentration to reduce the size of the sodium currents and increase potential for good voltage control (Colatsky, 1980). One could also examine the biophysical properties of single sodium channels in excised or cell attached patches from the cell body (Franciolini, 1986; Stühmer et al., 1987). Both of these would provide additional insight, but they also have the limitation of restricting analysis to a small population of channels in a restricted location (on or near cell body) that may not represent the full population of channels that define the firing properties of the neurons.

### 4.4 Hyperactive neural network

Our previous study demonstrates that iPSC-derived neurons form functionally active neural networks that contain GABAergic and glutamatergic synaptic connections (Xie et al., 2018), allowing us to evaluate the effect the K1270T mutation on the network activity of the neuronal cultures. Our results reveal an increase in excitatory drive relative to inhibitory drive in excitatory neurons.

There are no significant differences in the (E-I)/(E+I) index on the basis of sPSC frequency between the two lines within the same isogenic pairs. Therefore, it is likely that alteration in the index based on charge transfer is associate with that on PSC amplitude. However, how a mutation in Nav leads to increased PSC amplitude is still unclear. It is likely that homeostatic regulation in genes that encode post-synaptic receptors is involved. Future studies will include examination of expression of glutamate receptors such as NMDA and AMPA receptors and GABA receptors such as GABA_A_ receptors etc.

Hyperexcitable neural networks are implicated in epileptic disorders (Shao et al., 2019). Therefore, our iPSC-derived neuronal cultures provide insights into the underlying mechanism of how an epilepsy-associated *SCN1A* mutation gives rise to increased probability of seizures in patients. To understand the effect of the mutation on the spontaneous network activity and network synchrony more broadly, our future studies will use multi-electrode arrays that allow for mechanistic study and anti-epileptic drug screening in a high-throughput manner (Tidball and Parent, 2016).

### 4.5 Developmental maturation

Using our neuronal differentiation protocol that includes plating iPSC-derived neural progenitors onto astroglial feeder layers, the large majority of neurons fire multiple action potentials (Schutte et al., 2018; Xie et al., 2018). The derived neurons not only have sodium currents larger than 1 nA in average and fire multiple action potentials at 7-8 Hz comparable to other iPSC studies (Liu et al., 2013; Sun et al., 2016), but they also receive glutamatergic and GABAergic synaptic input within 3 weeks of plating. However, their electrophysiological properties are less mature than neurons expected in an adult brain, based on lower firing frequency, larger input resistance, and depolarized RMP. Single-cell transcriptome analysis reveals development of interneurons at D54 *in vitro* recapitulates those in human fetal cortex at 100 dpc (Close et al., 2017). While we cannot rule out the possibility that the K1270T mutation causes a developmental delay in neuronal development in culture, it seems unlikely to explain our results of reduced firing frequency and action potential AP in inhibitory neurons associated with the mutation because 1) there was no consistent change in RMP or input resistance and 2) the excitatory neurons of the mutated control and the patient lines were able to fire APs in similar or higher frequency compared to their controls. However future studies could evaluate effects of the mutation on further maturation in long-term cultures.

### 4.6 Off-target mutation

Sequences of the top five off-target sites from CRISPR/Cas9 editing, including *SCN7A*, *FLII*, *SCN2A*, *SCN3A* and *SCN9A* were examined. The corrected patient line did not have off-target mutations in the candidate genes (Supplementary Figure 1A). While the mutated control line did not have off-target mutations in the other genes, it harbored one in *SCN7A*. It is very unlikely that this mutation affected the results reported based on the following considerations. First, *SCN7A* encodes the Na_x_ sodium channel in heart, uterus, skeletal muscle, and peripheral nervous system (Goldin et al., 2000) and RT-PCR analysis demonstrated that it is not expressed in our iPSC-derived neurons of the mutated control line (Supplementary Figure 1B). Secondly, the channel is insensitive to changes in membrane potential and TTX, a voltage-gated sodium channel blocker (Hiyama et al., 2002). Isolated sodium currents in iPSC-derived neurons at D21-24 post plating were completely eliminated in the presence of 1 *µ*M TTX (Supplementary Figure 1C). Third, while it is possible that the off-target mutation in *SCN7A* could trigger homeostatic regulation by up- or down-regulating *SCN1A* expression, this is unlikely as the alterations in sodium currents/evoked firing in the homozygous mutated control line were in the same direction and twice as large as the heterozygous patient line that did not undergo editing process.

### 4.7 Comparison with the previous knock-in fly model

The reduced excitability observed specifically in iPSC-derived inhibitory neurons is consistent with the decreased excitability specifically in inhibitory neurons in the brains of K1270T GEFS+ knock-in *Drosophila* (Sun et al., 2012). Since both flies and patients exhibit heat-induced seizures, this provides additional support to the hypothesis that reduced firing specifically in inhibitory neurons is critical to the disease phenotype in humans. However, the cellular defect associated with reduced excitability is distinct between the models demonstrating two independent mechanisms that result in a similar phenotype. In the iPSC-derived neurons there is a significant decrease in the amplitude of the transient currents that are the dominant form of sodium current in these neurons. In flies the reduced firing is associated with a hyperpolarized shift in the deactivation threshold of the large persistent sodium currents characteristic of inhibitory neurons in the adult fly brain. It is possible that modifying the persistent currents in iPSCs by shifting their voltage dependence could moderate the deficits associated with a reduced transient sodium current.

In addition, an increase in excitability was observed in excitatory neurons of the patient line, consistent with the higher firing rate in excitatory neurons in the GEFS+ flies (Schutte et al., 2016). The alterations in fly are seen predominantly at high temperature, so it is possible that elevated temperatures will reveal additional or more severe alterations in sodium currents in iPSC-derived neurons, as the patient sibling from whom the skin fibroblast was collected showed more severe seizure phenotypes of febrile seizures compared to afebrile seizures (Abou-Khalil et al., 2001).

### 4.8 Summary

Overall the CRISPR/Cas9-based disease modeling strategy with iPSC-derived neurons can validate clinically relevant phenotypes and reveal the sufficiency and necessity of *SCN1A* mutations for specific phenotypes in the human genetic background. Furthermore, a library of patient iPSCs with a variety of *SCN1A* mutations built using this approach, in combination with a high-throughput drug screening method such as multi-electrode array (MEA), will facilitate the development of personalized anti-epileptic medicine (Du and Parent, 2015; Odawara et al., 2016; Parent and Anderson, 2015).

## 5. Supplemental information

**Supplementary Table 1.**
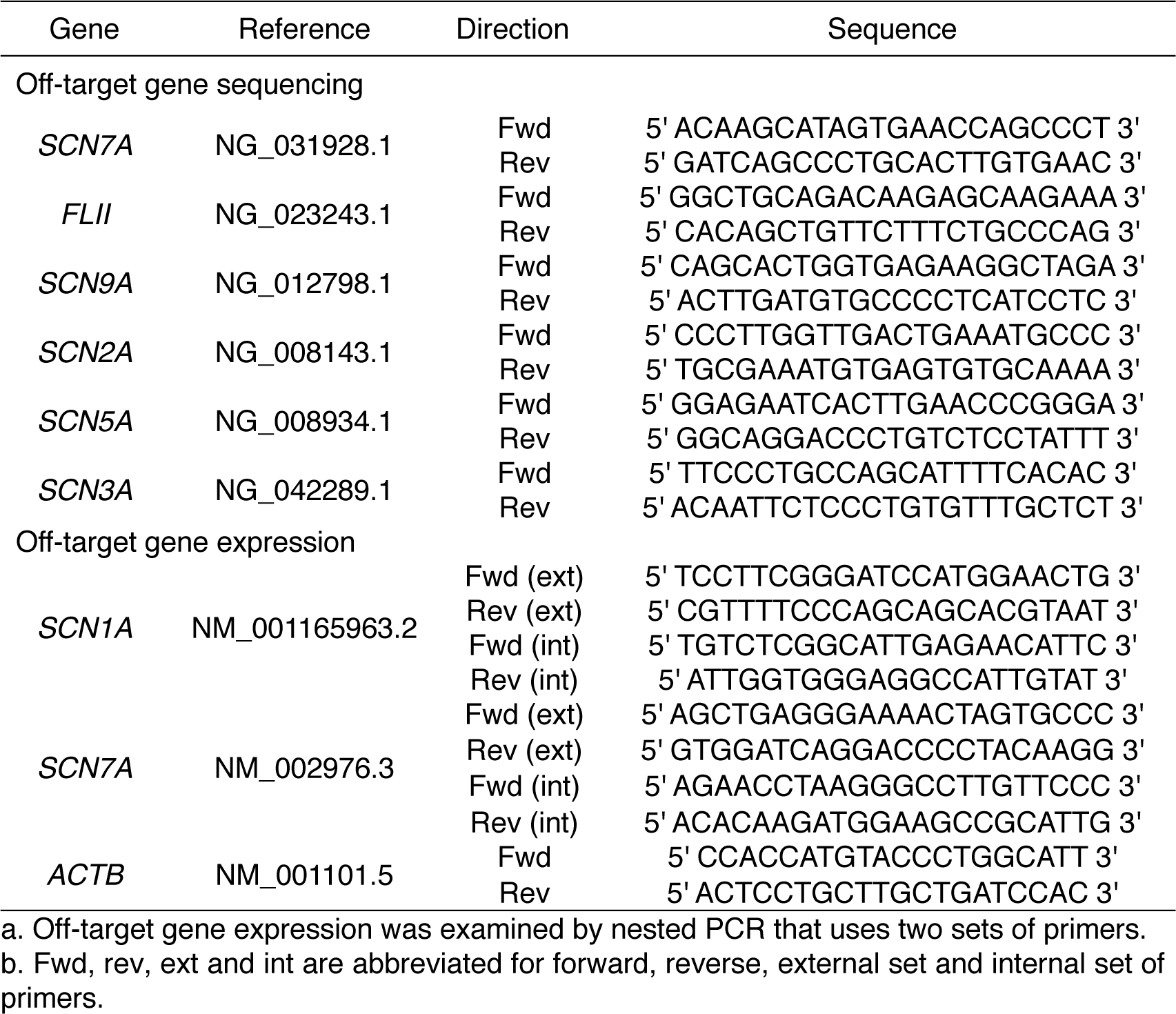
Primers for off-target gene sequencing and expression

**Supplementary Figure 1 related to Figure 1.**
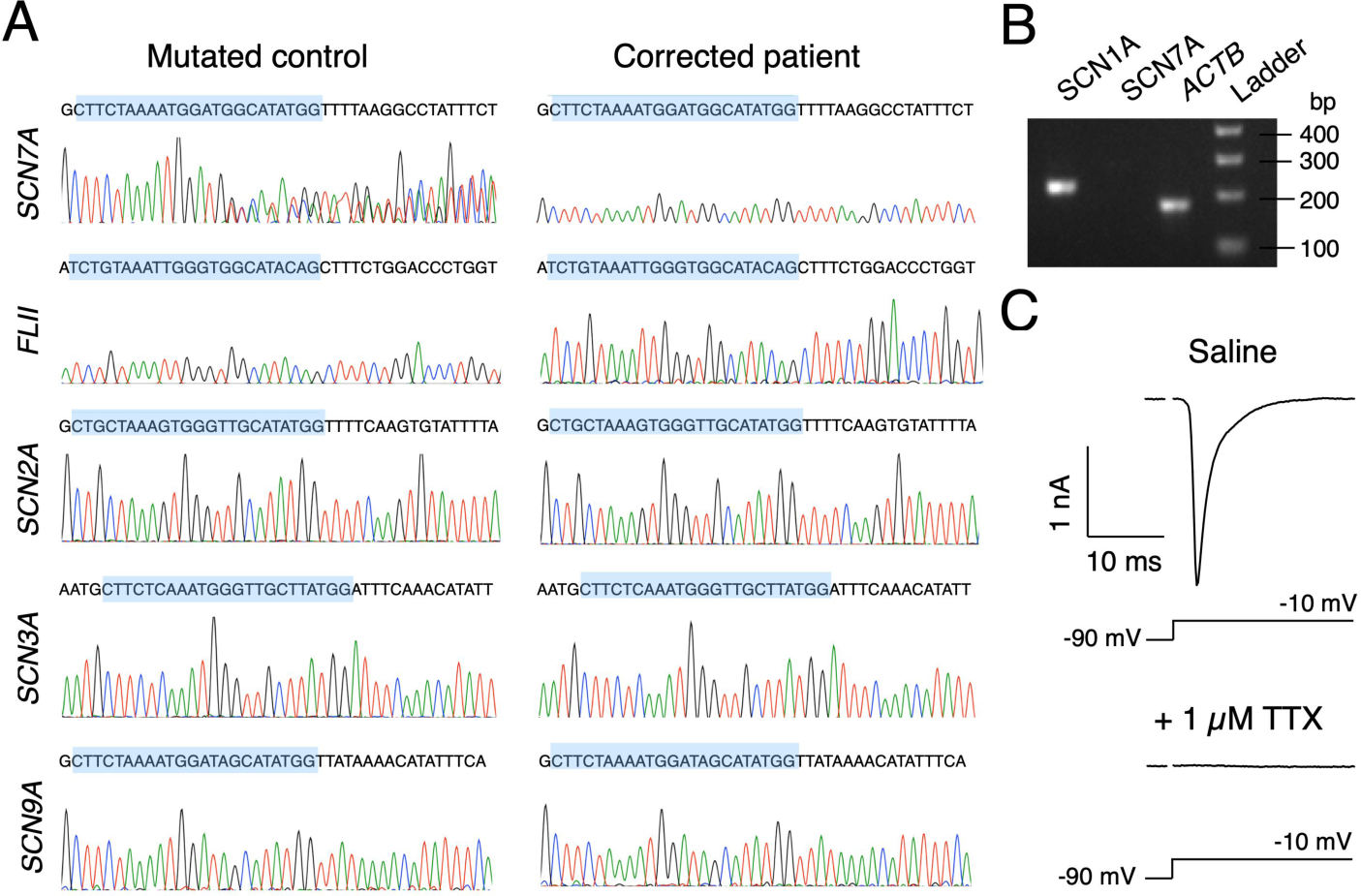
(A) Sequences of the top five predicted off-target sites in *SCN7A, FLII, SCN2A, SCN3A and SCN9A*. Off-target mutation was identified in the *SCN7A* gene of the mutated control line. Regions highlighted in blue are the sites predicted to be modified by non-specific CRISPR/Cas9 editing. (B) Expression of *SCN1A, SCN7A* and the housekeeping gene *ACTB* in the derived neurons of the mutated control line at D21 post plating. Expected band sizes of *SCN1A, SCN7A* and *ACTB* are 220, 283 and 177 bp. (C) Isolated sodium current recording in the derived neurons of the control line at D54 post plating. Sodium currents were completely eliminated in the presence of 1 μM TTX.

**Supplementary Figure 2 related to Figure 3.**
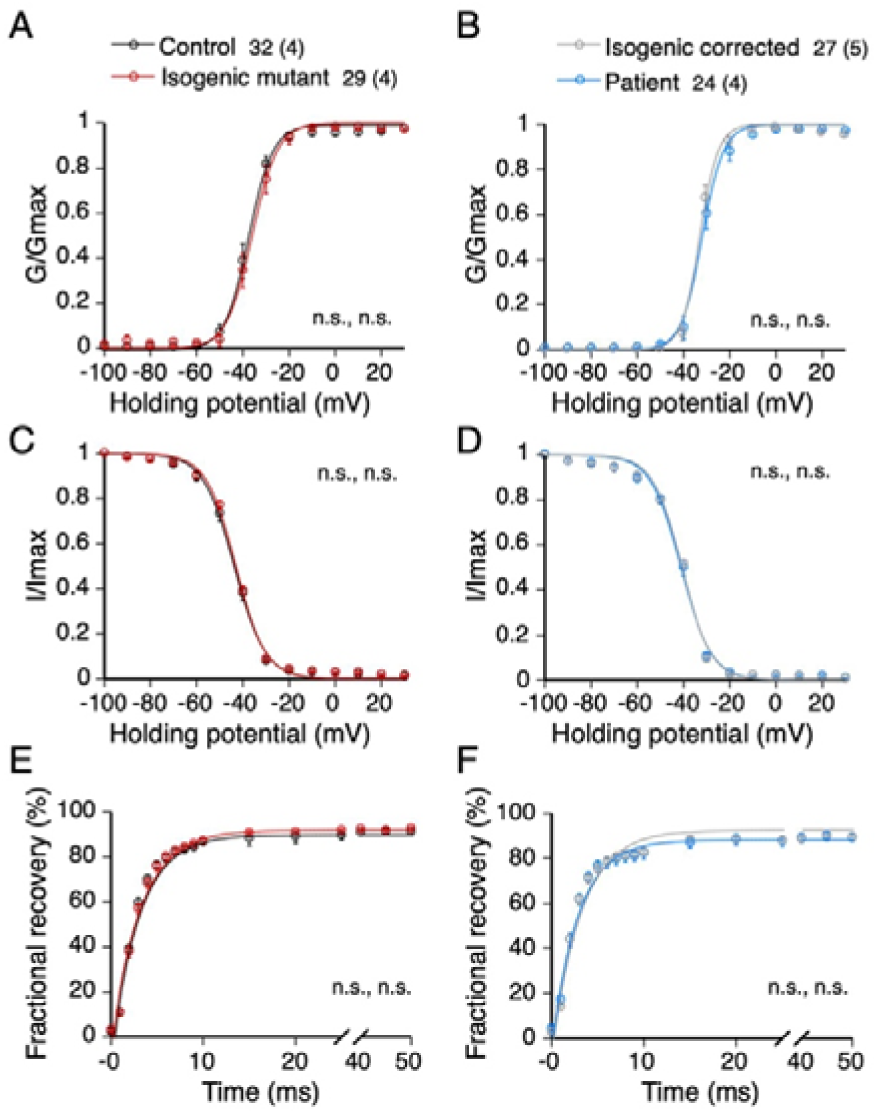
Properties of Nav in inhibitory neurons. (A-B) Conductance vs voltage curves of activation in inhibitory neurons of the control vs the mutated control lines, and the corrected patient vs the patient lines respectively. (C-D) Current vs voltage curves of steady-state inactivation in inhibitory neurons of isogenic pairs. (E-F) Recovery of inactivation in both isogenic pairs. Data presented as mean ± s.e.m with total number of neurons and number of individual platings in parentheses evaluated in each line. Two-way ANOVA and *post hoc* Sidak’s analysis.

**Supplementary Figure 3 related to Figure 4.**
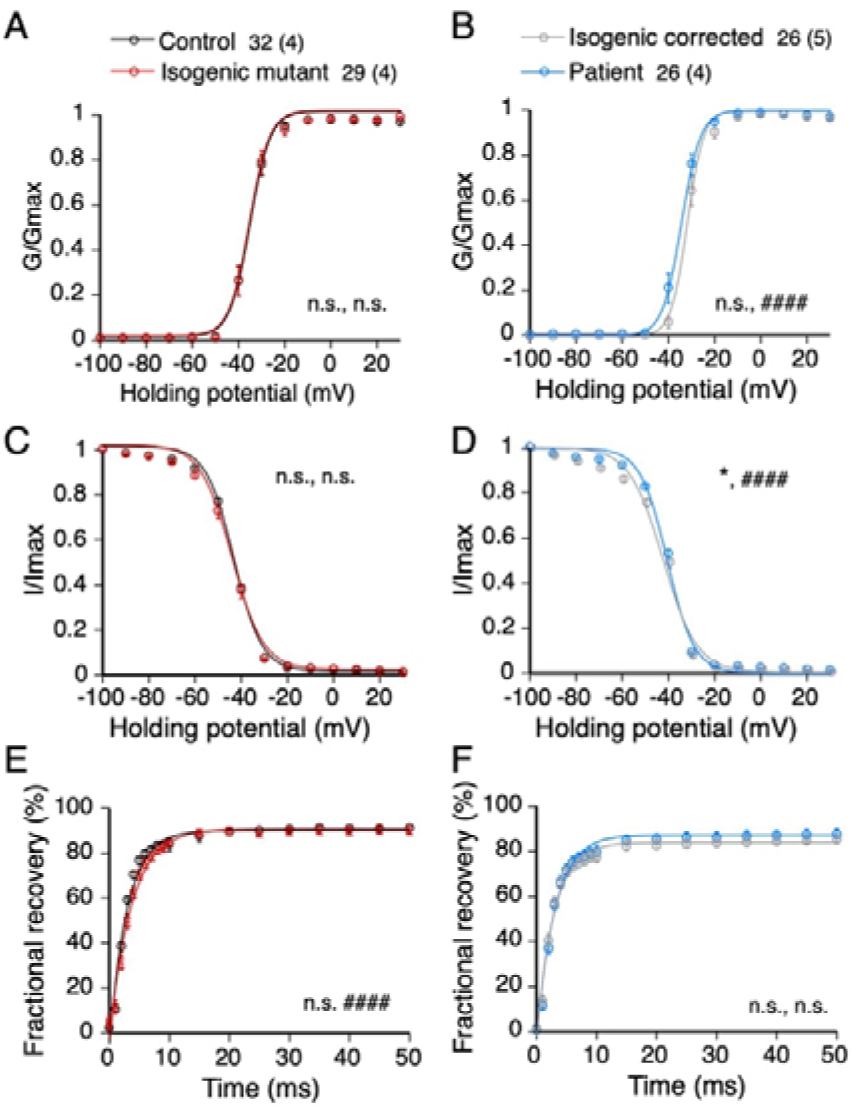
Properties of Nav in excitatory neurons. (A-B) Conductance vs voltage curves of activation in excitatory neurons of the control vs the mutated control lines, and the corrected patient vs the patient lines respectively. (C-D) Current vs voltage curves of steady-state inactivation in excitatory neurons of isogenic pairs. (E-F) Recovery of inactivation in both isogenic pairs. Data presented as mean ± s.e.m with total number of neurons and number of individual platings in parentheses. * and # represent significant differences in genotypes and interaction within an isogenic pair respectively, two-way ANOVA and *post hoc* Sidak’s analysis. One and four symbols denote p values < 0.05, and 0.0001 respectively.

**Supplementary Figure 4 related to Figure 5.**
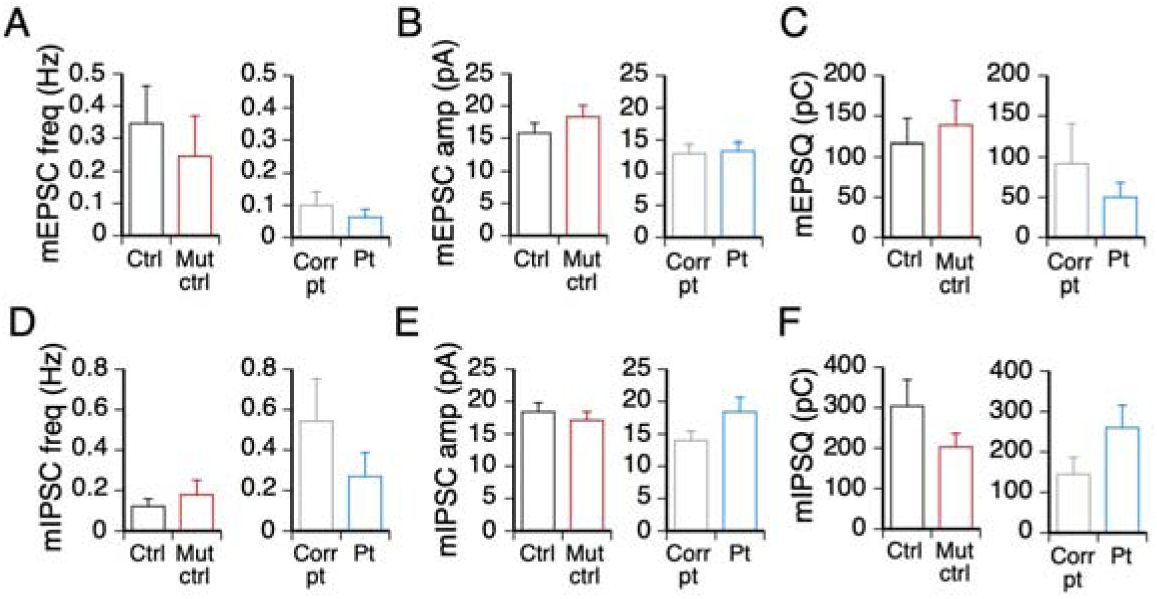
Miniature EPSCs IPSCs received by excitatory neurons were unaltered by the K1270T mutation. (A-C) The frequency, amplitude and charge transfer of mEPSCs in the control (ctrl) vs the mutated control (mut ctrl), and the corrected patient (corr pt) vs the patient (pt) lines. (D-F) Same as A-C, but for mIPSCs. Data reported as mean + s.e.m. The total number of neurons and number of individual platings in parentheses for all four lines are ctrl: 9 (5), mut ctrl: 7 (3), corr pt:10 (4), and pt:10 (3).

## 6. Acknowledgement

This work was supported by the National Institutes of Health grants R01 NS083009 (D.K.O.), R01 GHD059967 grant (P.H.S.), and R01 NS078289 (K.C.E.) and a California Institute for Regenerative Medicine Bridges to Stem Cell Research grant CIRM-EDUC2-08383 (C.M.R.). We would also like to thank Alexander E. Stover for generating the iPSCs, Priscilla Figueroa, Noor Osman and Daniel R. Benavides for help on astroglial cultures, Sara E. Konopelski for help on screening of clones, Karla Soto Sauza for help on immunostaining and cell counting and Longwen Huang for input on the manuscript.

## 7. Author contributions

The laboratory of K.C.E. provided skin fibroblasts from the unaffected sibling and the patient sibling from the GEFS+ family. The laboratory of P.H.S. reprogrammed the fibroblasts into iPSCs. O.S.S. and M.A.S. designed the CRISPR/Cas9, and O.S.S., Y.X., N.N.N. and C.M.R. performed screening of positive clones. C.M.R., N.N.N., O.S.S. and Y.X. prepared astroglial cultures. Y.X., N.N.N. and C.M.R differentiated iPSCs into neurons. Y.X. and N.N.N. performed electrophysiological recording. Y.X., N.N.N. and C.M.R analyzed the electrophysiology data. Y.X. and N.N.N. performed immunostaining, imaging and cell counting. Y.X. wrote the manuscript, N.N.N., O.S.S., K.C.E, P.H.S., M.A.S. and D.K.O. edited the manuscript.

